# Modelling illustrates that genomic selection provides new opportunities for intercrop breeding

**DOI:** 10.1101/2020.09.11.292912

**Authors:** Jon Bančič, Christian Werner, Chris Gaynor, Gregor Gorjanc, Damaris Odeny, Henry F. Ojulong, Ian K. Dawson, Steve Hoad, John M. Hickey

## Abstract

Intercrop breeding programs using genomic selection can produce faster genetic gain than intercrop breeding programs using phenotypic selection. Intercropping is an agricultural practice in which two or more component crops are grown together. It can lead to enhanced soil structure and fertility, improved weed suppression, and better control of pests and diseases. Especially in subsistence agriculture, intercropping has great potential to optimise farming and increase profitability. However, breeding for intercrop varieties is complex as it requires simultaneous improvement of two or more component crops that combine well in the field. We hypothesize that genomic selection can significantly simplify and accelerate the process of breeding crops for intercropping. Therefore, we used stochastic simulation to compare four different intercrop breeding programs implementing genomic selection and an intercrop breeding program entirely based on phenotypic selection. We assumed three different levels of genetic correlation between monocrop grain yield and intercrop grain yield to investigate how the different breeding strategies are impacted by this factor. We found that all four simulated breeding programs using genomic selection produced significantly more intercrop genetic gain than the phenotypic selection program regardless of the genetic correlation with monocrop yield. We suggest a genomic selection strategy which combines monocrop and intercrop trait information to predict general intercropping ability to increase selection accuracy in early stages of a breeding program and to minimize the generation interval.

## 1 Introduction

Intercropping is an agricultural practice in which two or more component crops are grown together (Vandermeer 1989). A common combination is a cereal with a legume, such as maize with beans in Latin America (Zimmermann 1996), and finger millet and pigeon pea in India (Dass and Sudhishri 2010). Intercropping can lead to enhanced soil structure and fertility, the conservation of soil moisture, improved weed suppression, and better control of pests and diseases, enabling greater yields and higher profitability (Brooker et al. 2015; Litrico and Violle 2015). It also allows for simultaneous cultivation of crops with different nutritional profiles, which can contribute to improving diets (Dawson et al. 2019a) and to increasing the stability and resilience of food systems (Himmelstein et al. 2017; Raseduzzaman & Jensen, 2017). Due to these characteristics, intercropping has great potential to optimise farming, especially in subsistence agricultural systems, which has recently led to an increased interest in the development and evaluation of efficient intercrop production (Dawson et al. 2019b).

Despite the potential benefits of intercropping, intercrop breeding has received only very little attention to date, with varieties specifically bred for intercrop production being unavailable (Brooker et al. 2015; Litrico and Violle 2015). This lack of attention is due to two major reasons:

i. In advanced economies, major global crop species are predominantly grown as monocrops (Leff et al. 2004) and the majority of breeding programs are focused on generating varieties adapted to monocrop production (Acquaah 2012).
ii. Intercrop breeding is more complex than monocrop breeding. Breeding for intercrop production requires the optimization of two or more component crops simultaneously (Wright 1985; Francis 1981); intercrop varieties ideally exhibit both a high *per se* performance and combine well with the other component crops (Davis and Woolley 1993).

As a result, the literature on intercrop breeding methodology is rare (Hill, 1996; Wright, 1985; Hamblin et al. 1976) and almost no progress in approaches has been made over the last few decades. The crop varieties currently used for intercropping have typically been bred for monocrop production, and most often their performance in intercropping has not even been evaluated in advance (Brooker et al. 2015), strongly restricting the potential benefits of this practice.

Genomic selection offers many opportunities to address the complexity of intercrop breeding programs and aid the simultaneous improvement of two or more component crops that combine well in the field. Genomic selection uses associations between genome-wide markers and phenotypic performance to predict the value of genotypes based on their genomic markers (Hickey et al. 2017; Lorenz et al. 2011; Meuwissen et al. 2001). In the context of an intercrop breeding program, genomic selection could be used in several ways to increase the rate of genetic gain:

i. Selection accuracy can be increased for individual performance and combined performance of the component crops in an intercrop.
ii. The generation interval can be reduced, since new crossing parents can be selected based on their genomic predicted values as soon as they are genotyped.
iii. Selection intensity can be increased, since thousands of potential intercrop combinations could be evaluated without testing all of them in the field.

We hypothesize that genomic selection can significantly simplify and accelerate the process of breeding crops for intercropping. To test our hypothesis, we used stochastic simulation to compare four different intercrop breeding programs implementing genomic selection and an intercrop breeding program using phenotypic selection only. We assumed three different levels of genetic correlation between monocrop grain yield and intercrop grain yield to investigate how the different breeding strategies are impacted by this factor. All four breeding programs using genomic selection produced significantly more intercrop genetic gain than the phenotypic selection program, regardless of the genetic correlation. We suggest a genomic selection strategy which combines monocrop and intercrop trait information to predict general intercropping ability to increase selection accuracy in early stages of a breeding program and to minimize the generation interval.

## 2 Material and Methods

Stochastic simulations were used to evaluate the potential of genomic selection for intercrop breeding. We compared four different intercrop breeding programs implementing genomic selection and an intercrop breeding program using phenotypic selection for long-term efficacy for maximizing intercrop grain yield. Below, we have subdivided the material and methods into three sections that describe first the simulation of the founder genotype population; second, the simulation of the recent (burn-in) breeding phase using a phenotypic selection breeding program; and third, the simulation of the future breeding phase to compare four different genomic selection breeding programs to the phenotypic selection breeding program. These topics are briefly reviewed below before detail is provided.

### Simulation of the founder genotype population

i. Genome simulation: a genome sequence was simulated for two hypothetical component crops in an intercrop production.
ii. Simulation of founder genotypes: the simulated genome sequences were used to generate a base population of 100 founder genotypes for each of the two component crops.
iii. Simulation of genetic values: for each of the two component crops, two traits were simulated, representing monocrop grain yield and intercrop grain yield. Genetic values for the two traits were calculated by summing the additive genetic effects for both traits at 10,000 quantitative trait nucleotides (QTN) and three different genetic correlations (0.4, 0.7, 0.9) were simulated.
iv. Simulation of phenotypes: phenotypes were simulated for monocrop grain yield and intercrop grain yield. Phenotypes representing monocrop grain yield were generated by adding random error to the genetic values for monocrop grain yield. Phenotypes representing intercrop grain yield were generated by adding random error to the mean genetic values for intercrop grain yield of two genotypes from both component crops.

### Simulation of the recent (burn-in) breeding phase

A phenotypic selection breeding program was simulated for 20 years (burn-in) to provide a common starting point for the comparison of the different intercrop breeding programs during the future breeding phase.

### Simulation of the future breeding phase

Four different genomic selection breeding programs were simulated and compared to the phenotypic selection breeding program for an additional 20 years of future breeding. The four genomic selection breeding programs included three variations of a Conventional genomic selection breeding program and a Grid genomic selection breeding program.

### 2.1 Simulation of the founder genotype population

#### 2.1.1 Genome simulation

Genome sequences were simulated for two hypothetical component crops used in intercropping. For modelling purposes, the two crops’ genomes were assumed to have the same characteristics. Each genome sequence consisted of 10 chromosome pairs. Each chromosome had a genetic length of 1.43 Morgans and a physical length of 8×10^8^ base pairs. The chromosome sequences were generated using the Markovian coalescent simulator (MaCS, Chen et al. 2009) implemented in AlphaSimR (Gaynor et al. 2020; R Core Team 2019). Recombination rate was derived as the ratio between genetic length and physical length (i.e., 1.43 Morgans/8×10^8^ base pairs = 1.8×10^−9^ per base pair). The per-site mutation rate was set to 2×10^−9^ per base pair. Effective population size was set to 50, with linear piecewise increases up to 32,000 at 100,000 generations ago, as described by Gaynor et al. (2017).

#### 2.1.2 Simulation of founder genotypes

The simulated genome sequences were used to generate a base population of 100 founder genotypes in Hardy-Weinberg equilibrium, for each of the two component crop species. These genotypes were formed by randomly sampling 10 chromosome pairs per genotype. A set of 1,000 bi-allelic QTNs and 2,000 single nucleotide polymorphisms (SNP) were randomly selected along each chromosome. This was done to simulate the structure of a quantitative trait that was controlled by 10,000 QTN and a SNP marker array with 20,000 genome-wide SNP markers. The founder genotypes were converted to doubled haploids (DH) and served as initial parents in the burn-in phase.

#### 2.1.3 Simulation of genetic values

For each of the two component crops, two traits were simulated:

i. Monocrop grain yield, representing the yield of a genotype under monocrop production.
ii. Intercrop grain yield, representing the total yield of two genotypes, each from one of the two component crops, under intercrop production.

Genetic values for the two traits were calculated by summing the additive genetic effects for both traits across all 10,000 QTN. Additive effects were sampled from a standard normal distribution and scaled to obtain an additive variance of 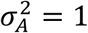 in the founder population, as described in detail in the vignette of the AlphaSimR package (Gaynor et al. 2020).

Three different genetic correlations (0.4, 0.7 and 0.9) were simulated to represent different degrees of genotype-by-cropping interaction (Davis and Woolley 1993).

#### 2.1.4 Simulation of phenotypes

Phenotypes were simulated for monocrop grain yield and for intercrop grain yield. Phenotypes for monocrop grain yield were generated by adding random error to the genetic values for monocrop grain yield. The random error was sampled from a normal distribution with mean zero and error variance 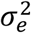, defined by the target level of heritability at each stage of a breeding program. Entry-mean values for narrow-sense heritability (*h*^2^) in the founder population were set to 0.1 in the doubled haploid stage and 0.33 in the preliminary yield trial stage. Narrow-sense heritabilities in later breeding stages increased as a result of an increased number of replicates (*r*) per genotype (**Tab. 1** and **Tab. 2**).

**Tab. 1.** Summary of *per stage* parameters and annual operational costs for four ‘medium’ breeding programs. Cost is set at approximately US $500K. The cost breakdown is shown for the Phenotypic selection breeding program (Pheno) and the three Conventional genomic selection breeding programs.

| Year | Stage | Reps | $h^2$ | Pheno | Baseline-GS | PYT-GS | DH-GS |
| --- | --- | --- | --- | --- | --- | --- | --- |
| 1 | Cross |  |  | 100 x 50 DH | 100 x 47 DH | 100 x 47 DH | 40 x 80 DH |
| 2 | DH | 1 | 0.10 | 5,000 | 4,700 | 4,700 | 3,200 |
| 3 | PYT | 4 | 0.33 | 500 | 500 | 500 | 500 |
| 4 | GIA 1 | 4 | 0.33 | 50<sup>1</sup> | 50<sup>1</sup> | 50<sup>1</sup> | 50<sup>1</sup> |
| 5 | GIA 2 | 24 | 0.50 | 13<sup>2</sup> | 13<sup>2</sup> | 13<sup>2</sup> | 13<sup>2</sup> |
| 6 | SIA 1 | 16 | 0.67 | 9<sup>*</sup> | 9<sup>*</sup> | 9<sup>*</sup> | 9<sup>*</sup> |
| 7 | SIA 2 | 32 | 0.80 | 3<sup>*</sup> | 3<sup>*</sup> | 3<sup>*</sup> | 3<sup>*</sup> |
| Cost (US\$) | | | | 493,200 | 492,200 | 492,200 | 495,200 |

Baseline-GS, the Baseline genomic selection breeding program; PYT-GS, the Preliminary yield trial genomic selection breeding program; DH-GS, the Doubled haploid genomic selection breeding program; Reps, the effective number of replications (i.e. locations); $h^2$ , narrow-sense heritability; DH, the doubled haploid stage; PYT, the preliminary yield trial stage; GIA 1 and 2, the general intercropping ability stages 1 and 2; SIA 1 and 2, the specific intercropping ability stages 1 and 2. <sup>1</sup> denotes testing with one probe variety; <sup>2</sup> denotes testing with three probe varieties; \* number of specific intercrop combinations.

**Tab. 2.** Summary of *per stage* parameters and annual operational costs for the ‘medium’ Grid genomic selection breeding program (Grid-GS). Cost is set at approximately US $500K.

| Year | Stage | Reps | $h^2$ | Grid-GS |
| --- | --- | --- | --- | --- |
| 1 | Cross |  |  | 40 x 90 |
| 2 | DH | 1 | 0.10 | 3,600 |
| 3 | Grid | 1 | 0.10 | 900/250,000* |
| 4 | SIA1 | 16 | 0.67 | 50* |
| 5 | SIA2 | 32 | 0.80 | 8* |
| Cost (US\$) | | | | 493,800 |

Reps, the effective number of replications (i.e. locations); $h^2$ , narrow-sense heritability; DH, the doubled haploid stage; Grid, the Grid stage; SIA 1 and 2, the specific intercropping ability stages 1 and 2; \* number of specific intercrop combinations.

Phenotypes for intercrop grain yield were generated by adding random error to the mean genetic values for intercrop grain yield of two genotypes from the two component crops. The following equation was used to calculate intercrop grain yield:

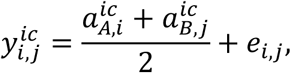

where 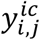 is the intercrop grain yield, 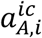 is the genetic value for intercrop grain yield of genotype i from component crop A, 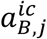 is the genetic value for intercrop grain yield of genotype j from component crop B, and *e*_*i,j*_ is the random error. The random error for intercrop grain yield was also sampled from a normal distribution with mean zero and error variance defined by the target level of heritability at each stage of a breeding program.

Narrow-sense heritabilities for monocrop grain yield and intercrop grain yield were calculated as:

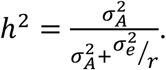

### 2.2 Simulation of the recent (burn-in) breeding phase

The phenotypic selection breeding program was simulated for 20 years (burn-in) to provide a common starting point for the comparison of the five intercrop breeding programs during the future breeding phase.

In brief, the four key features of the phenotypic selection breeding program (**Fig. 1**) were:

**Fig. 1:**
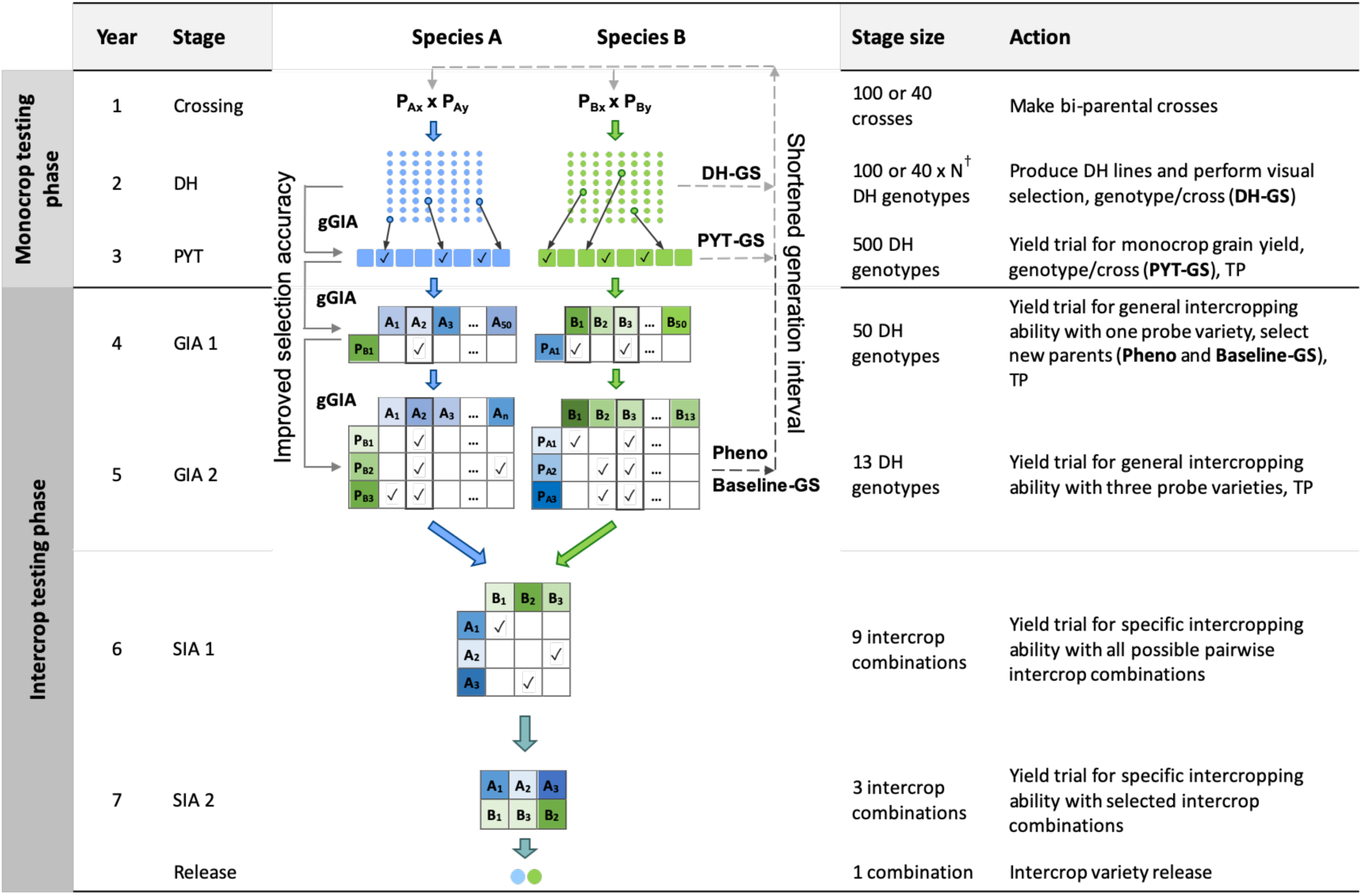
Schematic overview of the Phenotypic selection breeding program (Pheno) and the three Conventional genomic selection breeding programs. Baseline-GS, the Baseline genomic selection breeding program; PYT-GS, the Preliminary yield trial genomic selection breeding program; DH-GS, the Doubled haploid genomic selection breeding program; gGIA, genomic-predicted general intercropping ability; TP, denotes stages in which genotypic and/or phenotypic records are collected; DH, the doubled haploid stage; PYT, the preliminary yield trial stage; GIA 1 and 2, the general intercropping ability stages 1 and 2; SIA 1 and 2, the specific intercropping ability stages 1 and 2. Solid line with arrow represents increased selection accuracy based on gGIAs and dashed line with arrow represents shortened generation interval. †The number of DH lines per cross (N) differs for each breeding program to maintain equal operating costs.

i. A crossing block of 80 DH lines was used to develop 100 bi-parental populations each year for each of the two component crops.
ii. New DH lines were developed from each bi-parental cross.
iii. A two-year monocrop testing phase, in which monocrop grain yield was evaluated for each of the two component crop species, respectively.
iv. A four-year intercrop testing phase, in which intercrop grain yield was evaluated for each intercrop combination of two genotypes from the two component crops. New parents were selected in the second year of the intercrop testing phase, giving a generation interval of five years.

The time from crossing to the release of a pair of component crop varieties for intercrop production was seven years.

In what follows, the four key features of the phenotypic selection breeding program are explained in more detail. Simulation parameters, including heritability, the number of trial replications, and the number of tested genotypes at each stage of the breeding program, are shown in Table 1. In the context of the phenotypic selection breeding program for intercropping varieties, we introduce the following terms:

i. Probe variety: represents a genotype of one of the two component crops that has good intercropping ability with genotypes from the other component crop. It is comparable to a tester in a hybrid breeding program, which is used to evaluate the general combining ability of genotypes from one heterotic pool with another heterotic pool.
ii. General intercropping ability (GIA): the average intercrop grain yield of a genotype from one component crop grown with genotypes from the other component crop. It is evaluated using one or several probe genotypes from the other component crop.
iii. Specific intercropping ability (SIA): the intercrop grain yield of a specific intercrop combination of two genotypes from the two component crops.

#### Crossing block (year 1)

Each year, for each crop, a crossing block of 80 DH lines was used to produce 100 bi-parental crosses (**Tab. 1**). Parental combinations were chosen at random from all 3,160 possible pairwise combinations.

#### Development of doubled haploids (year 2)

From each bi-parental cross, 50 DH lines were produced for each of the two component crops. The resulting 5,000 DH lines per crop were advanced to the monocrop testing phase and tested in the same year.

#### Monocrop testing phase (years 2 and 3)

The monocrop testing phase spanned two years. Performance was evaluated as monocrop grain yield. Monocrop testing included the doubled haploid stage (DH stage, year 2) and the preliminary yield trial stage (PYT stage, year 3). In the DH stage, seed was increased and phenotypic selection was based on single plants within each bi-parental cross, to ensure there was variation in the later stages. In the PYT stage, phenotypic selection was based on multi-location trial plots (**Tab. 1**).

#### Intercrop testing phase (year 4 to 7)

The intercrop testing phase spanned four years. Performance was evaluated as intercrop grain yield, i.e., the total yield from both simultaneously grown genotypes of the two component crops. The intercrop testing phase included two general intercropping ability testing stages (GIA1, year 4; and GIA2, year 5) and two specific intercropping ability testing stages (SIA1, year 6; and SIA2, year 7).

In the GIA1 stage, phenotypic selection was based on general intercropping ability in a yield trial with one probe variety. In the GIA2 stage, phenotypic selection was based on general intercropping ability using three probe varieties. Each year, the 4 best performing genotypes from the GIA2 stage replaced the probe varieties from the previous year. New parents were selected in the GIA1 stage. Each year, the 20 best performing genotypes from the GIA1 stage were used to replace the 20 oldest parents in the crossing block. Hence, every genotype stayed in the crossing block for four crossing cycles. The generation interval was five years.

In the SIA1 stage, all possible pairwise combinations of the selected lines were tested in a yield trial. In the SIA2 stage, the best combinations from the SIA1 stage were tested in a multi-location trial (**Tab. 1**). The highest yielding intercrop combination was then released as an intercrop variety combination.

### 2.3 Simulation of the future breeding phase

The future breeding phase was used to evaluate the phenotypic selection breeding program and the four genomic selection breeding programs for an additional 20 years of breeding. The genomic selection breeding programs included three variations of a Conventional genomic selection breeding program and a Grid-GS breeding program. The three Conventional genomic selection breeding programs replaced phenotypic selection by genomic selection at different stages of the phenotypic selection breeding program (**Fig. 1**). They comprised a Baseline-GS, a PYT-GS and a DH-GS breeding program. The Grid-GS breeding program reorganized the phenotypic selection breeding program to enable the evaluation of a greater number of specific intercrop combinations using genomic selection. Details on all programs are given below.

In order to obtain approximately equal annual operating costs, the number of doubled haploids per bi-parental cross was reduced in the genomic selection breeding programs to compensate for additional costs due to genotyping. Table 1 shows the resources used for the phenotypic selection breeding program and the three Conventional genomic selection breeding programs. Table 2 shows the resources used for the Grid-GS breeding program. Estimated applied costs in calculations were $20 per monocrop test plot, $50 per intercrop test plot, $35 for producing a doubled haploid line and $20 for producing a single genotype by array genotyping.

#### 2.3.1 The Baseline genomic selection breeding program (Baseline-GS)

In the Baseline-GS breeding program, genomic selection was used to replace phenotypic selection in the PYT stage and in the GIA1 stage. Each year, the best 20 genotypes from the GIA1 stage were selected as new parents using genomic selection to replace the oldest 20 parents in the crossing block. As for the phenotypic selection breeding program, the generation interval was 5 years.

#### 2.3.2 The Preliminary yield trial genomic selection breeding program (PYT-GS)

In the PYT-GS breeding program, genomic selection was used to replace phenotypic selection in the PYT stage and in the GIA1 stage. Each year, the best 80 genotypes from the PYT stage and last year’s crossing block were selected as new parents using genomic selection.

#### 2.3.3 The Doubled haploid genomic selection breeding program (DH-GS)

In the DH-GS breeding program, genomic selection was used to replace phenotypic selection in the DH stage, the PYT stage and the GIA1 stage. Each year, 80 genotypes from the DH stage were selected as new parents to replace the entire last year’s crossing block. As preliminary results showed a rapid decrease in genetic variance when the best 80 genotypes were selected, we implemented a maximum avoidance crossing scheme using genomic selection to reduce the rate of decrease (Kimura and Crow 1963).

#### 2.3.4 Genomic selection model for the three Conventional genomic selection breeding programs

The three Conventional genomic selection breeding programs used a multivariate ridge regression genomic selection model (RR-BLUP) to obtain genomic predictions of general intercropping abilities (gGIA) for each component crop separately.

In this model, monocrop grain yield from the PYT stage and mean intercrop grain yield with one or three probes respectively from the GIA1 and the GIA2 stage were fitted simultaneously. Genomic predictions of general intercropping ability can be directly calculated using intercrop grain yield from the GIA1 and the GIA2 stage as phenotypic information. In addition, the multivariate model uses information on monocrop grain yield, which was included as a correlated trait.

The following model was used:

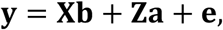

expanded in matrix form as:

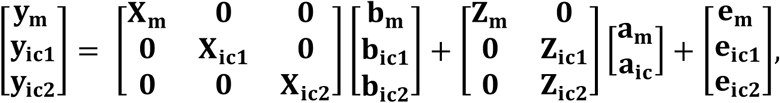

where **y**_**m**_, **y**_**ic1**_ and **y**_**ic2**_ respectively denote the vectors of monocrop grain yield from the PYT stage, and mean intercrop grain yield with one or three probes from the GIA1 and the GIA2 stage; **b**_**m**_, **b**_**ic1**_and **b**_**ic2**_ respectively denote the vectors for the fixed effects of year and stage for PYT, GIA1, and GIA2; **a**_**m**_ and **a**_**ic**_ respectively denote the vectors of the marker effects for monocrop grain yield and intercrop grain yield; **X**_**m**_, **X**_**ic1**_, **X**_**ic2**_, **Z**_**m**_, **Z**_**ic1**_ and **Z**_**ic2**_ denote the corresponding incidence matrices; and **e**_**m**_, **e**_**ic1**_and **e**_**ic2**_ denote the corresponding vectors of residuals.

Additive genetic (**G**) and residual (**R**) variance-covariance matrices were:

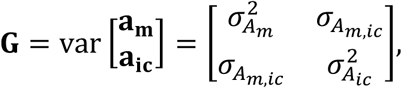

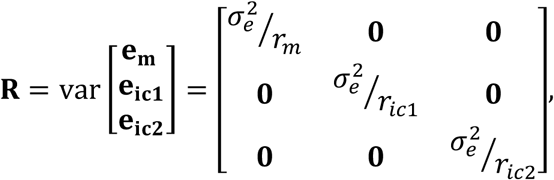

where 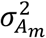 and 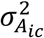 respectively denote the additive genetic variances for monocrop grain yield and intercrop grain yield, and 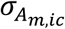 denotes the additive genetic covariance between the two traits; and 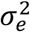 denotes the residual variance. **R** modelled heterogeneous residual variances by weighting 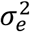 for the effective number of replications (*r*) in a particular stage (**Tab. 2**). To reduce computation time, additive genetic variances were assumed known and calculated each year using the true additive genetic effects.

The initial training population at the start of the future breeding phase consisted of all genotypes from the PYT stage of the last five years of the burn-in phase. This training population consisted of 2,500 genotypes and 2,739 phenotypic records from the PYT, the GIA1 and the GIA2 stages. In every year of the future breeding phase, 500 new genotypes from the PYT stage were added to the training population, as well as 500, 50 and 13 new phenotypic records from the PYT, the GIA1 and the GIA2 stages, respectively. The training population was updated using a 5-year sliding window approach, in which it always contained the most recent 5 years of training data.

#### 2.3.5 Grid genomic selection breeding program (Grid-GS)

The Grid genomic selection breeding program reorganized the phenotypic selection breeding program to enable the evaluation of a greater number of specific intercrop combinations using genomic selection (**Fig. 2**). To achieve this, the PYT, the GIA1 and GIA2 stages were replaced by a single grid stage. The reorganized program design also tested an increased number of specific intercrop combinations at the SIA1 and the SIA2 stages **(Tab. 2)**.

**Fig. 2:**
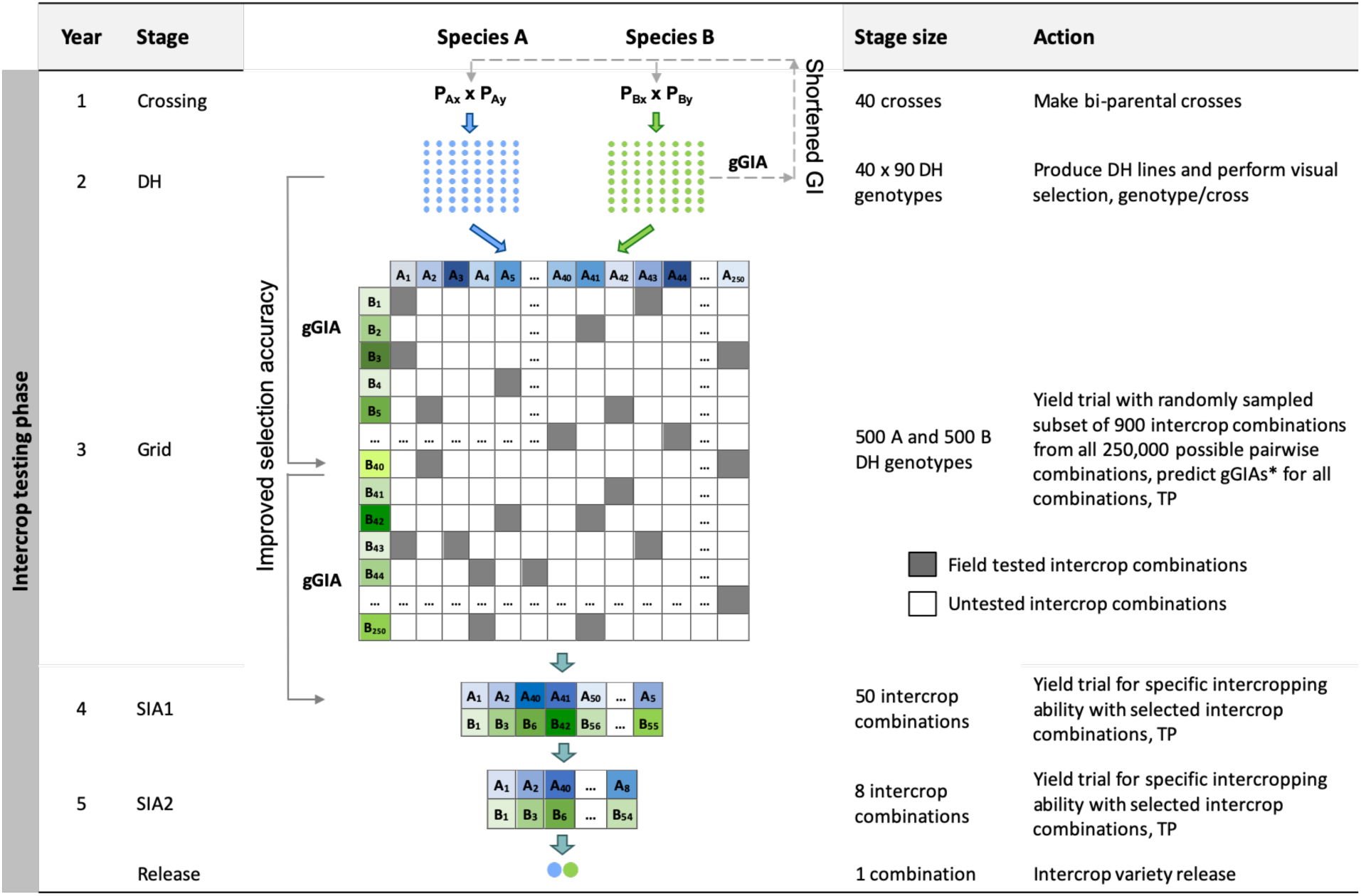
Schematic overview of the Grid genomic selection breeding program (Grid-GS). gGIA, genomic-predicted general intercropping ability; TP, denotes stages in which genotypic and/or phenotypic records are collected; DH, the doubled haploid stage; Grid, the Grid stage; SIA 1 and 2, the specific intercropping ability stages 1 and 2. Solid line with arrow represents increased selection accuracy based on gGIAs and dashed line with arrow represents shortened generation interval.

The Grid stage involved field testing of 900 intercrop combinations. At first, genomic prediction of general intercropping ability (gGIA) was used at the DH stage to select the best 500 DH lines from each component crop. From all 250,000 possible intercrop combinations between the 500 DH lines from each component crop, 900 were randomly sampled for field testing at a single location (**Tab. 2**).

Genomic predictions of general intercropping ability were calculated for all 250,000 intercrop combinations using the following equation:

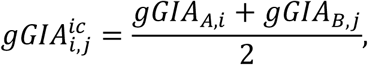

where 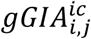 is the mean genomic-predicted general intercropping ability; and *gGIA*_*A,i*_ and *gGIA*_*B,i*_ are respectively the genomic-predicted general intercropping abilities of the *i*-th and *j*-th genotypes of component crops A and B. The best 50 predicted intercrop combinations were then advanced to the SIA1 stage (compared to nine intercrop combinations in the phenotypic selection breeding program and the three Conventional genomic selection breeding programs).

Each year, 80 genotypes from the DH stage were selected as new parents to replace the entire last year’s crossing block. As preliminary results showed a rapid decrease in genetic variance when the best 80 genotypes were selected, we implemented a maximum avoidance crossing scheme with genomic selection to reduce the rate of decrease (Kimura and Crow 1963). The generation interval was two years and the total length of the breeding program from initial crosses to the release of the intercrop variety pair was five years.

#### 2.3.6 Genomic selection model for the Grid-GS breeding program

Genomic predictions of general intercropping ability were calculated using a ridge regression model (RR-BLUP) which predicted marker effects for both component crops simultaneously based on intercrop grain yield.

The following model was used:

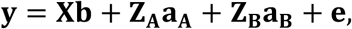

expanded in matrix form as:

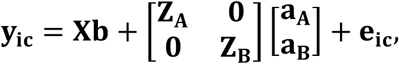

where **y**_**ic**_ denotes the vector of intercrop grain yield from the Grid, SIA1 and SIA2 stages; **b** denotes the vector for fixed effects of year and stage for Grid, SIA1, and SIA2; **a**_**A**_ and **a**_**B**_ respectively denote the vectors of the marker effects for intercrop grain yield in component crops A and B; **X, Z**_**A**_ and **Z**_**B**_ denote the incidence matrices; and **e**_**ic**_ denotes the vector of residual effects.

The residual (**R**) variance-covariance matrix modelled heterogeneous residual variances by weighting 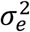 for the effective number of replications (*r*) in a particular stage (**Tab. 4**), as described for the genomic selection model used in the Conventional genomic selection breeding programs. To reduce computation time, additive genetic variances were assumed known and calculated each year using the true additive genetic effects.

To initialise the training population, the Grid stage was already simulated during the last five years of the burn-in breeding phase. For each of the five years, the best 23 genotypes at the PYT stage for each component crop were selected based on their genomic-predicted general intercropping abilities. These selected genotypes were then used to generate all 529 possible intercrop combinations, which were then tested in the field. At the beginning of the future breeding phase, the initial training population thus consisted of 115 genotypes from each component crop and 2,645 intercrop grain yield phenotypes (5 x 529 different intercrop combinations). In every year of the future breeding phase, 500 new genotypes from each component crop were added to the training population, as well as 900, 50 and 8 intercrop grain yield records respectively from the Grid, SIA1 and SIA2 stages. The training population was updated using a 5-year sliding window approach, in which the training population always contained the most recent five years of data. This training population contained a total of 2,500 genotypes and 4,790 phenotypic records.

### 2.4 Comparison of the intercrop breeding program designs

The performance of the five intercrop breeding programs (the four using genomic selection and the phenotype-alone comparison) was evaluated by measuring the mean intercrop genetic value over time in the DH stage of both component crops as follows:

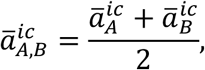

with 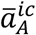 and 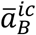 being the mean intercrop genetic values of the genotypes in the DH stage from component crops A and B, respectively. Mean intercrop genetic values of the two component crops were centered at 0 for the last year of the burn-in breeding phase. Intercrop genetic variance was measured as variance of the mean intercrop genetic values. Direct comparisons between breeding program designs for intercrop genetic gain and intercrop genetic variance were reported as ratios. These were calculated by performing a paired t-test (Welch) on log-transformed values from the 30 simulation replicates; the log-transformed differences from the t-test were then back-transformed to obtain ratios (**Tab. S5**; Gaynor et al. 2017).

Prediction accuracy was evaluated as the correlation coefficient between the true and predicted performance at the DH stage. In the phenotypic selection breeding program, the phenotype served as a predictor of intercropping ability. Prediction accuracy was evaluated as the correlation between the phenotypic value (i.e., monocrop grain yield) and true intercrop genetic value. In all four genomic selection breeding programs, prediction accuracy was measured as the correlation between the genomic-predicted general intercropping ability and the true intercrop genetic value of the doubled haploids.

Comparisons of the five breeding programs were done under three different levels of annual operating budget: (i) a ‘large’ budget (US $1M); (ii) a ‘medium’ budget (US $500K); and (iii) a ‘small’ budget (US $250K). Since the results for all our breeding programs showed similar rankings across these budgets, the methods presented above and the results presented below are described only for the medium budget scenario (**Tab. 1 and Tab. 2**). The parameters applied for breeding program designs and the results of the simulations at other budget levels are presented in the Supplementary Materials (**Tab. S1, S2, S3, and S4; Fig. S1 and S2**).

## 3 Results

Our results show that intercrop breeding programs using genomic selection can produce faster genetic gain than an intercrop breeding program using only phenotypic selection. All four breeding programs using genomic selection produced more intercrop genetic gain than the phenotypic selection breeding program, regardless of the genetic correlation between monocrop grain yield and intercrop grain yield. However, the three Conventional genomic selection breeding programs produced increasingly more genetic gain when the genetic correlation between monocrop grain yield and intercrop grain yield increased, while the Grid-GS breeding program produced slightly less genetic gain when the genetic correlation increased. The DH-GS breeding program always produced the most genetic gain among the three Conventional genomic selection breeding programs. Intercrop breeding using genomic selection also gave a faster reduction in genetic variance than intercrop breeding with phenotypic selection, regardless of the genetic correlation between monocrop yield and intercrop yield. Selection accuracy for intercropping ability was higher when genomic selection was compared to phenotypic selection. Selection accuracy in the three Conventional genomic selection breeding programs and the phenotypic selection breeding program increased when the genetic correlation between monocrop grain yield and intercrop grain yield increased, while selection accuracy in the Grid-GS breeding program was similar under different levels of genetic correlation. The general trends and rankings observed under the medium annual budget were representative of the trends observed under low and high annual budgets. Our findings are discussed in more detail below in terms of gain, genetic variance and selection accuracy.

### 3.1 Intercrop genetic gain

Intercrop breeding using genomic selection produced faster genetic gain than intercrop breeding with phenotypic selection. This is shown in Figure 3, which plots intercrop genetic gain as mean intercrop genetic value in the DH stage for the entire future breeding phase. The three panels show intercrop genetic gain for the five simulated breeding programs under different levels of genetic correlation between monocrop grain yield and intercrop grain yield. All four breeding programs using genomic selection produced significantly more intercrop genetic gain than the phenotypic selection program under all three levels of genetic correlation between monocrop yield and intercrop yield.

**Fig. 3.**
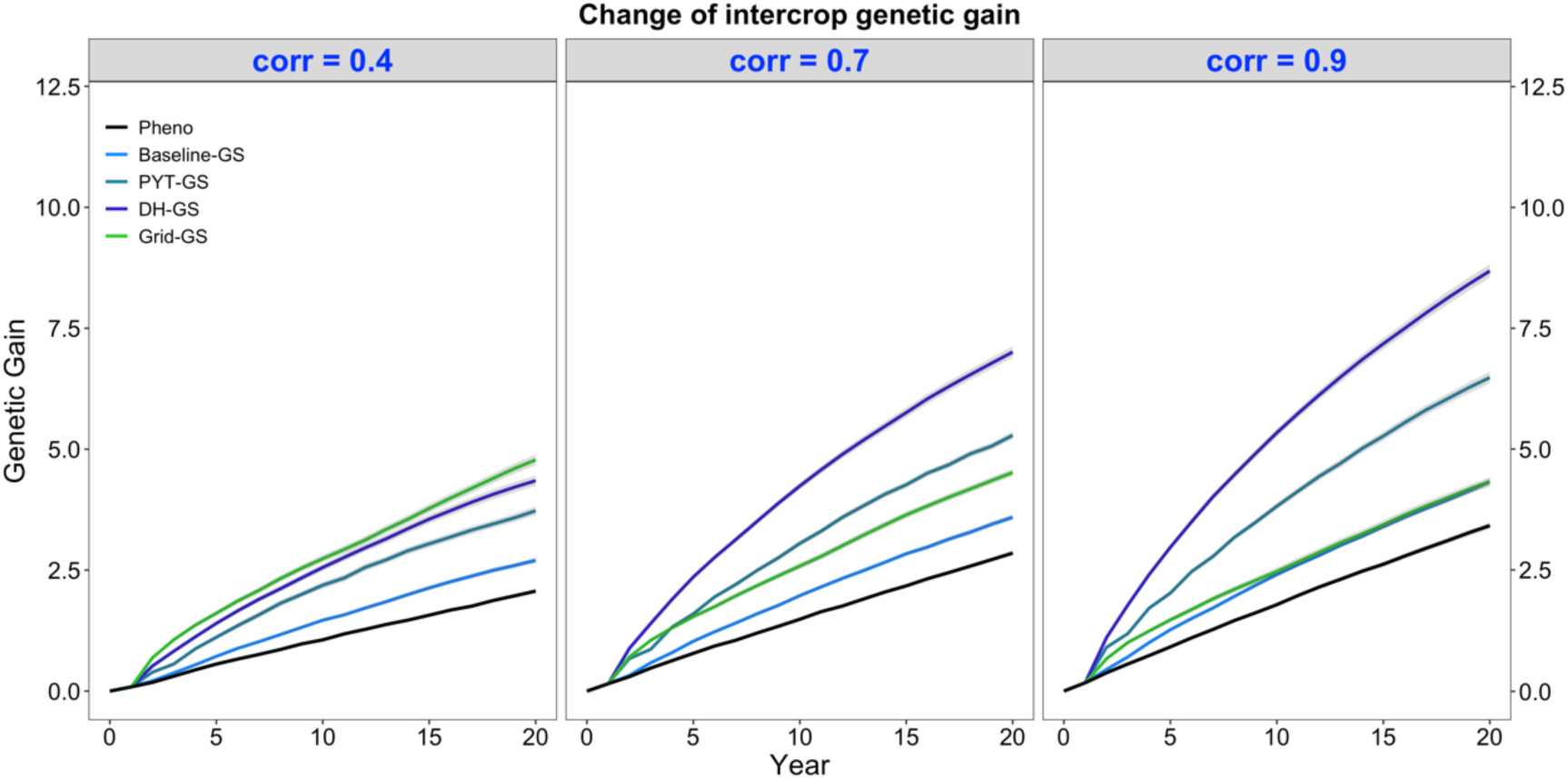
Intercrop genetic gain over time for five simulated breeding programs. Results are shown under genetic correlations of 0.4, 0.7 and 0.9. Simulations are based on a medium annual operating budget (approx. US $500K). Intercrop genetic gain is plotted as mean intercrop genetic value in the doubled haploid stage for the entire future breeding phase. The lines within each of the three panels represent the five breeding programs where each line represents mean genetic value for the 30 simulated replicates and the shadings show standard error bands. The black line represents the Phenotypic selection breeding program (Pheno), the blue-coloured lines represent the three Conventional genomic selection breeding programs (Baseline-GS, the Baseline genomic selection breeding program; PYT-GS, the Preliminary yield trial genomic selection breeding program; DH-GS, the Doubled haploid genomic selection breeding program) and the green-coloured line represents the Grid genomic selection breeding program (Grid-GS).

Figure 3 also shows that the three Conventional genomic selection breeding programs produced increasingly more genetic gain when the genetic correlation between monocrop grain yield and intercrop grain yield increased, while the Grid-GS breeding program produced slightly less genetic gain when the genetic correlation increased. As a result, the ranking of the four genomic selection breeding programs for genetic gain was dependent on the level of genetic correlation. When the genetic correlation was low (0.4), the Grid-GS breeding program produced the most genetic gain over time, closely followed by the DH-GS breeding program. Both breeding programs produced more than twice the genetic gain of the phenotypic selection breeding program. However, when the genetic correlation was high (0.9), the Grid-GS breeding program produced less genetic gain than all three Conventional genomic selection breeding programs. It generated 1.3 times the gain of the phenotypic selection breeding program, while the DH-GS breeding program produced 2.5 times the genetic gain of the phenotypic selection breeding program.

Figure 3 also shows that the DH-GS breeding program always produced the most genetic gain of the three Conventional genomic selection breeding programs, followed by PYT-GS and Baseline-GS breeding programs. The relative performance of the DH-GS breeding program compared to the other two Conventional genomic selection breeding programs increased when the genetic correlation between monocrop grain yield and intercrop grain yield increased. When the genetic correlation was low (0.4), the DH-GS breeding program generated 1.2 times the genetic gain of the PYT-GS breeding program and 1.6 the gain of the Baseline-GS breeding program. When the genetic correlation was high (0.9), it generated 1.3 times the genetic gain of the PYT-GS breeding program and twice the gain of the Baseline-GS breeding program.

All breeding programs produced more genetic gain when the annual operating budget was high (**Tab. S5; Fig S2a**) and less genetic gain when the annual operating costs were low (**Tab. S5; Fig S1a**). The general trends and rankings observed under the medium annual budget were representative of the trends observed under low and high annual budgets.

### 3.2 Intercrop genetic variance

Intercrop breeding using genomic selection gave a faster reduction in genetic variance than intercrop breeding with phenotypic selection. This is shown in Figure 4, which plots the genetic variance of the intercrop genetic values in the DH stage for the entire future breeding phase. All four breeding programs using genomic selection gave a faster reduction in genetic variance than the phenotypic selection breeding program under all three levels of genetic correlation between monocrop yield and intercrop yield.

**Fig. 4.**
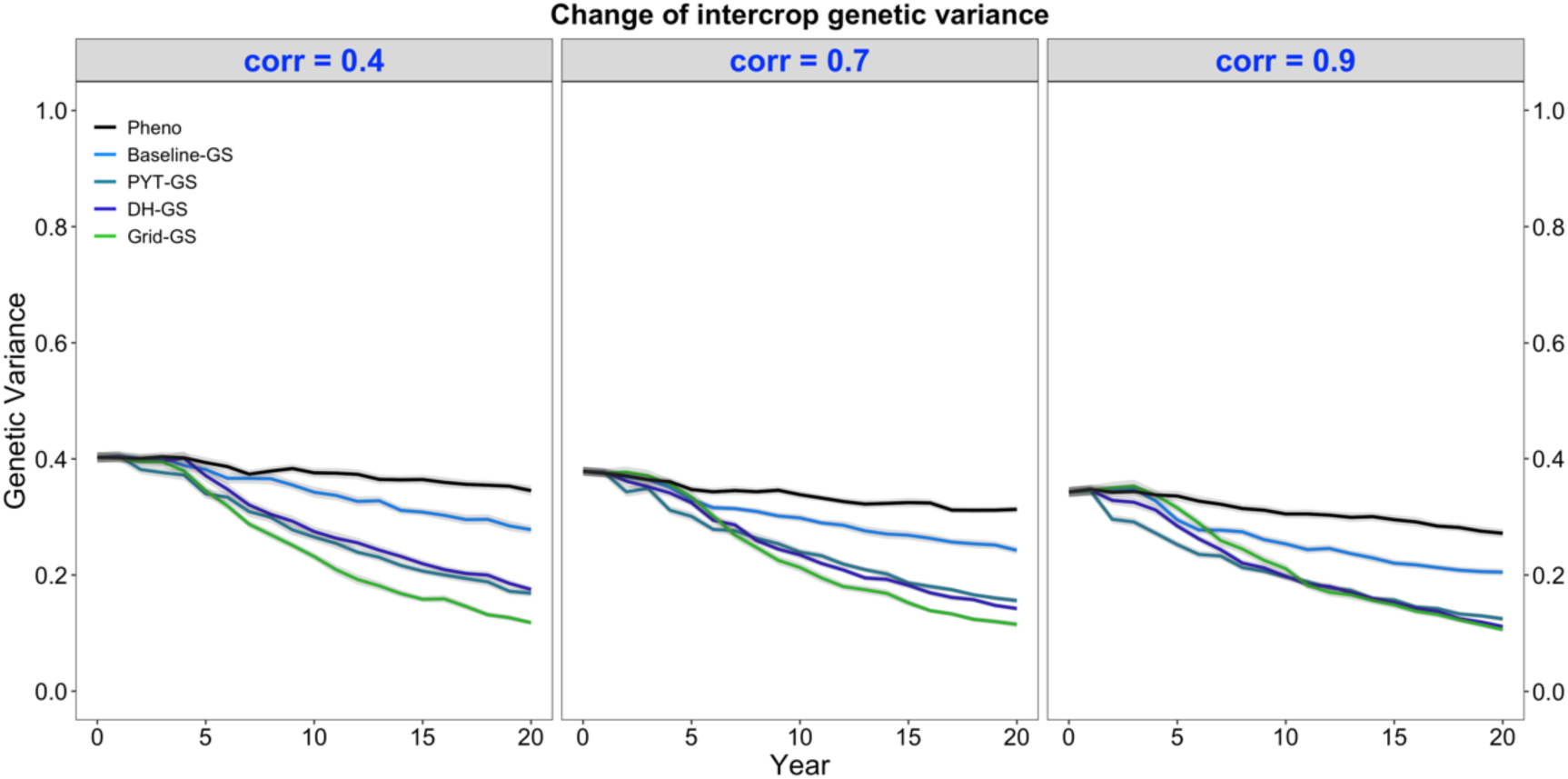
Intercrop genetic variance over time for five simulated breeding programs. Results are shown under genetic correlations of 0.4, 0.7 and 0.9. Simulations are based on a medium annual operating budget (approx. US $500K). Intercrop genetic variance is plotted as variance of intercrop genetic values in the doubled haploid stage for the entire future breeding phase. The lines within each of the three panels represent the five breeding programs where each line represents mean intercrop genetic variance for the 30 simulated replicates and the shadings show standard error bands. The black line represents the Phenotypic selection breeding program (Pheno), the blue-coloured lines represent the three Conventional genomic selection breeding programs (Baseline-GS, the Baseline genomic selection breeding program; PYT-GS, the Preliminary yield trial genomic selection breeding program; DH-GS, the Doubled haploid genomic selection breeding program) and the green-coloured line represents the Grid genomic selection breeding program (Grid-GS).

Figure 4 also shows that the Grid-GS breeding program gave the fastest reduction in genetic variance at the end of the future breeding phase under all three levels of genetic correlation. The Baseline-GS breeding program gave the slowest reduction in genetic variance among the four breeding programs using genomic selection. The DH-GS and the PYT-GS breeding programs always showed a similar reduction in genetic variance and ranked between the other two breeding programs using genomic selection. However, these two breeding programs became more similar to the Grid-GS breeding program as the genetic correlation increased. When the genetic correlation was high (0.9), the Grid-GS, the PYT-GS and the DH-GS breeding programs all showed a similar reduction in genetic variance at the end of the future breeding phase.

All breeding programs gave a faster reduction in genetic variance when the annual operating budget was high (**Fig S1b**) and a slower reduction in genetic variance when annual operating costs were low (**Fig S2b**). The general trends observed under the medium annual budget were representative of the trends observed under low and high annual budgets.

### 3.3 Genomic selection accuracy

Genomic selection for intercropping ability was more accurate than phenotypic selection for intercropping ability. This is shown in Figure 5, which plots the mean selection accuracy for general intercropping ability in the DH stage for the entire future breeding phase. All four breeding programs using genomic selection showed on average higher accuracy than the phenotypic selection breeding program under all three levels of correlation between monocrop yield and intercrop yield.

**Fig. 5.**
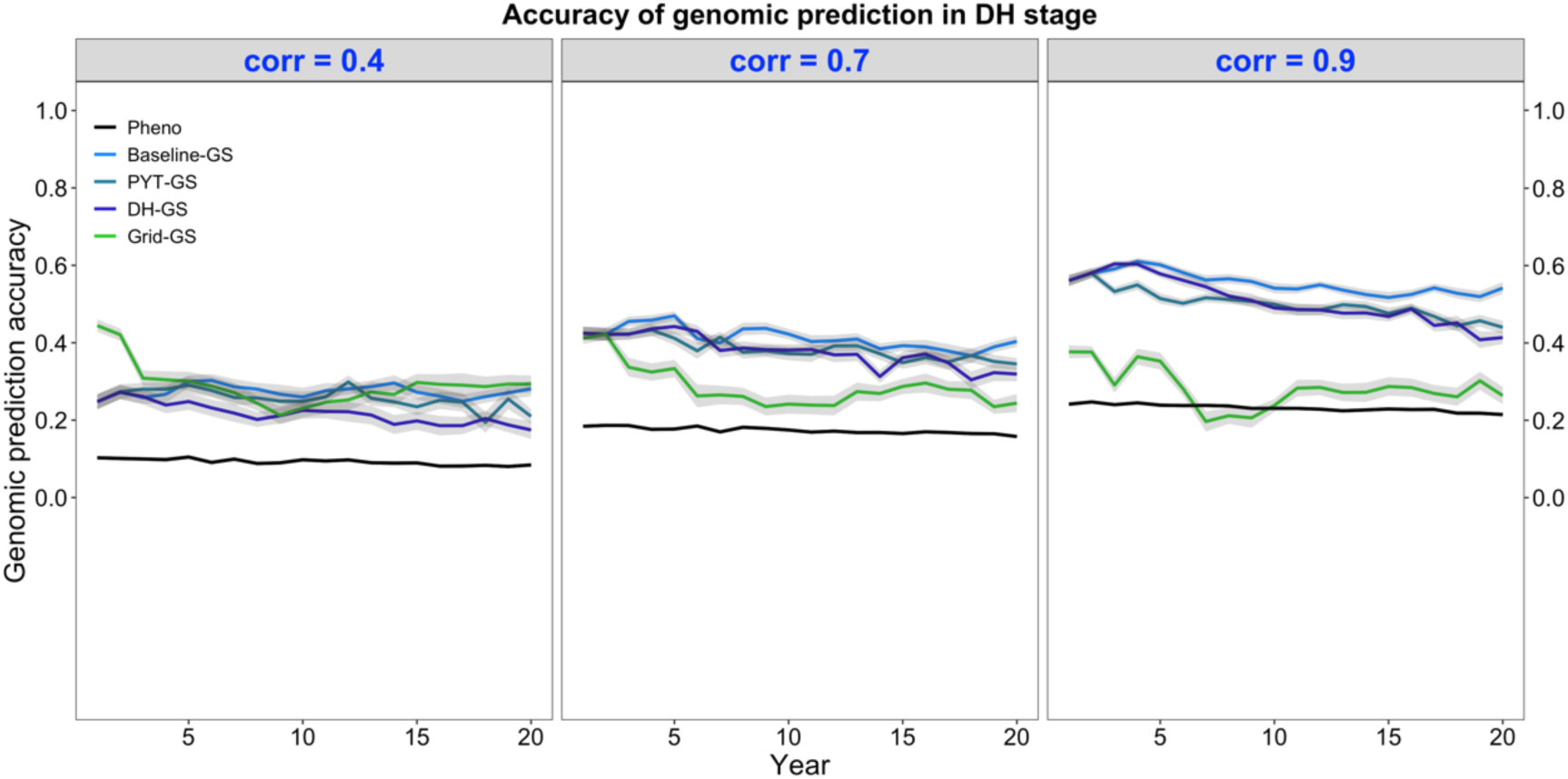
Genomic prediction accuracy over time for five simulated breeding programs. Results are shown under genetic correlations of 0.4, 0.7 and 0.9. Simulations are based on a medium annual operating budget (approx. US $500K). Genomic prediction accuracy is plotted as mean genomic-predicted general intercropping ability in the doubled haploid stage for the entire future breeding phase. The lines within each of the three panels represent the five breeding programs where each line represents mean genomic-predicted general intercropping ability for the 30 simulated replicates and the shadings show standard error bands. The black line represents the Phenotypic selection breeding program (Pheno), the blue-coloured lines represent the three Conventional genomic selection breeding programs (Baseline-GS, the Baseline genomic selection breeding program; PYT-GS, the Preliminary yield trial genomic selection breeding program; DH-GS, the Doubled haploid genomic selection breeding program) and the green-coloured line represents the Grid genomic selection breeding program (Grid-GS).

Figure 5 also shows that selection accuracy for intercropping ability in the three Conventional genomic selection breeding programs and the phenotypic selection breeding program increased when the genetic correlation between monocrop grain yield and intercrop grain yield increased. Selection accuracy in the Grid-GS breeding program, on the other hand, was on average similar under all three levels of genetic correlation. When the genetic correlation was low (0.4), all four breeding programs using genomic selection showed on average a relatively similar selection accuracy. However, when the genetic correlation was high (0.9), the three Conventional genomic selection breeding programs showed a significantly higher selection accuracy than the Grid-GS breeding program. During most of the future breeding phase, the selection accuracy of the Grid-GS breeding program was even lower than the selection accuracy of the phenotypic selection breeding program.

The four breeding programs using genomic selection showed a higher selection accuracy when the annual operating budget was high (**Tab. S5; Fig S1a**) and a lower selection accuracy when the annual operating costs were low (**Tab. S5; Fig S2a**). The general trends observed under the medium annual budget were representative of the trends observed under low and high annual budgets.

## 4 Discussion

High-performance intercrop production systems require more efficient intercrop breeding approaches that make use of advances in breeding (Dawson *et al*., 2019b). While it offers potential advantages, genomic selection also incurs additional costs, so it is necessary to understand the balance between benefits and costs. Stochastic simulations are becoming widely used to explore the efficiency of genomic selection in single species breeding only (e.g. Muleta et al. 2019; Gorjanc et al. 2018; Gaynor et al. 2017), but to our knowledge our use of simulations to explore genomic selection’s value for intercrop breeding is unique. Through simulations, we have shown that intercrop breeding programs using genomic selection can produce faster genetic gain than intercrop breeding programs which use phenotypic selection, working to a common cost basis that reflects the resources available for a mediumly-invested breeding initiative.

To discuss our results, we first examine the value of genomic selection to increase selection accuracy and reduce the generation interval in breeding crops for intercrop production. We also explain that maximising the rate of genetic gain using genomic selection can significantly increase genetic gain in the short term, but may impair long-term genetic gain due to rapid depletion of genetic variance. We then describe the value of strategies which reduce the loss of genetic variance to solve this problem, such as maximum avoidance crossing schemes or optimal contribution selection. We explain why the performance of the different genomic selection breeding programs was dependent on the genetic correlation between monocrop grain yield and intercrop grain yield, and we conclude that the DH-GS breeding program should be used in intercrop breeding unless the genetic correlation between the two traits is known to be low. We finish by discussing the major limitations of our simulations and explain why we believe that our results are still valid in the context of real-world intercrop breeding programs.

### 4.1 Genomic selection increases intercrop genetic gain

In phenotypic selection breeding programs, new crossing parents are usually selected after several years of intensive testing in multiple environments. This enables high selection accuracies but also results in long generations intervals, substantially restricting the rate of genetic gain. Replacing phenotypic selection by genomic selection increases selection accuracy in early testing stages and thereby allows for selection of new parents based on their genomic predicted performance as soon as they can be genotyped. Our observations showed that all the intercrop breeding programs using genomic selection that we tested produced faster genetic gain than the phenotypic selection breeding program. This was observed regardless of the genetic correlation between monocrop grain yield and intercrop grain yield, and under three operating budgets. We observed that the major drivers of increased genetic gain were both an increased selection accuracy in early selection stages and a reduction of the generation interval. Our results were consistent with those of Gaynor et al. (2017) who used stochastic simulations to evaluate genomic selection strategies in plant breeding programs for developing inbred lines. We refer the reader to this study for a more detailed analysis of the relationship between genetic gain, the generation interval and prediction accuracy. As a consequence of increased selection accuracy and the reduced generation interval, all four genomic selection breeding programs also showed a faster reduction in genetic variance over time compared to the phenotypic selection breeding program. We discuss particular features of our findings in more detail below.

#### 4.1.1 Genomic selection accelerates the reduction of intercrop genetic variance over time

We found that all intercrop breeding programs using genomic selection showed a faster reduction of genetic variance than the phenotypic selection breeding program. As genomic selection improved the conversion of genetic variance into genetic gain, the accelerated reduction of intercrop genetic variance was a direct outcome of the increased selection accuracy and the reduced generation interval. While maximising this conversion will significantly increase the rate of genetic gain in the short term, the long-term genetic gain may be impaired due to a rapid depletion of genetic variance. To solve this problem, maximum avoidance crossing schemes can be used, which maintain genetic variation by selecting the best genotypes within families while ensuring that each family equally contributes to the next generation (Kimura and Crow 1963). In this way, an over-representation of the top families in future generations is prevented.

We experimented with this approach by applying a maximum avoidance crossing scheme in the DH-GS and the Grid-GS breeding programs, as initial simulations using genomic truncation selection of new parents resulted in rapid exhaustion of genetic variance. The approach was successful, but a downside of maximum avoidance crossing schemes is that they require a closed population with a constant number of families and a minimum number of progeny per family to ensure the least related crosses are made each generation. While these requirements can be easily met within a simulation framework, practical application of a maximum avoidance crossing scheme may be more challenging, as breeders might introduce new genetic material to their breeding population, and not every crossing event might produce seed. Other, more complex, strategies might be more suitable to reduce the loss of genetic variation in real-world breeding programs, such as optimal contribution selection and crossing (Gorjanc et al. 2018; Akdemir & Sánchez, 2016; Sonesson et al. 2012; Meuwissen, 1997), and exploring these could be a feature of future work.

#### 4.1.2 The DH-GS breeding program produces the most genetic gain when the genetic correlation between monocrop yield and intercrop yield is medium to high

In our simulations the DH-GS breeding program produced approximately two times the genetic gain of the Grid-GS breeding program and approximately 2.5 times the genetic gain of the phenotypic selection breeding program when the genetic correlation between monocrop yield and intercrop yield was medium to high (0.7 and 0.9). The DH-GS scheme benefited from a short generation interval and an increased selection accuracy in the DH, PYT and GIA1 stages.

To obtain genomic predictions of general intercropping abilities for each component crop, the DH-GS breeding program used a multivariate genomic selection model which fitted monocrop grain yield and intercrop grain yield simultaneously. While phenotypic information on intercrop yield came from the GIA1 and the GIA2 stages, the multivariate model enabled us to extract additional information from monocrop yield phenotypes due to the genetic correlation between monocrop yield and intercrop yield. This additional information resulted in increased selection accuracy when the correlation was medium to high. While novel in the context of intercrop breeding, the use of correlated traits in multivariate genomic models is a well-known approach to improve prediction accuracy with wide application in plant and animal breeding (Jia & Jannink, 2012; Calus & Veerkamp, 2011).

The same multivariate genomic selection model was also used in the Baseline-GS and the PYT-GS breeding programs. These two breeding programs also outperformed the phenotypic selection breeding program regardless of the genetic correlation between monocrop yield and intercrop yield, but produced less genetic gain than the DH-GS breeding program. The PYT-GS breeding program benefited from an increased selection accuracy and a reduced generation interval compared to the phenotypic selection breeding program. The Baseline-GS breeding program did not reduce the generation interval. It was used to demonstrate the increase in selection accuracy when genomic selection is used compared to phenotypic selection.

#### 4.1.3 The Grid-GS breeding program is advantageous when genetic correlation is low

In our modelling the Grid-GS breeding program produced approximately 1.2 times the genetic gain of the DH-GS breeding program and approximately 2.3 times the genetic gain of the phenotypic selection breeding program when the genetic correlation between monocrop yield and intercrop yield was low (0.4). Our findings can be explained by the fact that the Grid-GS genomic selection model did not consider monocrop yield records, so it is unaffected by the genetic correlations between monocrop yield and intercrop yield, and prediction accuracies are similar under all correlations. When the genetic correlation was low, it therefore outperformed the DH-GS breeding program. However, under high genetic correlation, the training population size of the DH-GS breeding program was effectively increased by including phenotypic information from monocrop stages, while the training population in the Grid-GS stage was not affected by the level of genetic correlation. We hypothesize that a larger training population and a better sampling strategy of intercrop combinations in the Grid stage would further increase the predictive ability and performance of the Grid-GS breeding program.

#### 4.1.4 Unless the genetic correlations between monocrop and intercrop yield are known to be low, the DH-GS breeding program should be used

Unless the genetic correlation between monocrop yield and intercrop yield was low, the DH-GS breeding program produced the most genetic gain. Even when the genetic correlation was 0.4, it was only slightly outperformed by the Grid-GS breeding program. These results indicate that the DH-GS breeding program has great potential to improve breeding for intercrop production.

In practical intercrop breeding programs, the genetic correlation between monocrop traits and intercrop traits will most likely be unknown and can change over time. The estimation of these parameters is difficult and requires large and costly experimental designs (Hill, 1996; Wright, 1985; Hamblin et al. 1976). When data for a precise decision-making process is not available, a strategy is required that delivers consistent performance across the whole parameter space. The DH-GS breeding program achieved substantially higher genetic gains than the phenotypic selection breeding program under all simulated correlations. Hence, we recommend that it is suitable for prompt implementation without prior knowledge about the level of genetic correlation.

A further advantage of the DH-GS breeding program is that it is a relatively simple way to implement genomic selection on top of a phenotypic selection intercrop breeding program, as it only requires minor resource re-allocations to compensate for the extra cost of genotyping. The Grid-GS breeding program, on the other hand, requires extensive restructuring of the breeding program, which might be harder to realise, particularly in low- and middle-income countries with limited resources.

### 4.2 Limitations of applying stochastic simulations for intercrop breeding program design

Our simulations have revealed the value of applying genomic selection in intercrop breeding. However, they are based on various simplified assumptions and do not model the full complexity of an actual intercrop breeding program. In this section, we discuss the major limitations of our simulations and explain why we believe that our results remain valid for real-world intercrop breeding. In the below we will discuss in turn assumptions which impact genomic selection accuracy, assumptions about making crosses and seed production, assumptions about the complexity of the breeding goal and assumptions about the absence of genotype-by-genotype interaction between the two component crops.

#### 4.2.1 Assumptions which impact genomic selection accuracy

The intercrop prediction accuracies obtained in our simulations are likely to be higher than those realized under real-world conditions. Our simulations may be inflated because: the variance components provided to genomic models were estimated directly from the true simulation parameters; there were no genotyping or phenotyping errors in our data set; and we assumed additive gene action without epistasis and genotype-by-environment interaction. Although these factors may affect genomic selection accuracies in a practical breeding program, the magnitude of the differences between the best genomic selection breeding programs and the phenotypic selection breeding program we have observed lead us to believe that in a real-world situation the benefits of genomic selection would remain tangible.

#### 4.2.2 Assumptions about making crosses and seed production

To minimize complexity, in our simulated breeding programs we assumed no differences in flowering time between crossing parents and that all crosses produce sufficient amounts of seed for immediate next step implementation. In real-world breeding, differences in maturity between potential crossing parents might reduce the number of possible crosses, while some crosses may not immediately produce enough seed, with additional seed multiplication steps required that prolong the breeding process. If dealing with a self-pollinating and an outcrossing component crop simultaneously, these issues might present the most significant challenge (Hamblin and Zimmermann 1986). As Hamblin et al. (1976) indicate, two self-pollinating crops with large seed production may be the simplest case for intercrop breeding. Breeding programs that use either phenotypic or genomic selection would be similarly affected by these seed production issues. We thus assume that the relative performance of the different breeding programs would be similar under more realistic crossing scenarios.

#### 4.2.3 Assumptions about the complexity of the breeding goal

In our analysis, comparisons between breeding programs were based on a single quantitative trait representing intercrop grain yield. We also assumed that both component crops equally contributed to intercrop grain yield and its economic value. Real-world breeding programs, however, have to consider multiple quantitative and qualitative traits simultaneously to maximise agronomic performance. Furthermore, it is unlikely that both component crops produce comparable amounts of yield and that both component traits have a similar market value.

In fact, the contribution of each component crop to the total economic value of the combined product will depend on various factors. These include: the cultivation environment (i.e. biological, economic and cultural) and management practices (Francis 1981; Mead and Riley 1981); the *per se* yield potential and economic value of each component crop (Wright 1985; Francis 1981; Hamblin, Rowell, and Redden 1976); and the intended use of the products, especially whether for subsistence use or for market (Mead and Riley 1981).

In theory, a selection index could be developed to enable selection of the best intercrop combinations by combining several key traits and through considering the above factors. Selection indices allow assigning customized economic weights to the component crops, thereby optimising their individual yield gains to maximize the market value of the combined crop product. In real-world breeding programs, estimation of the relative (economic) weights for traits of interest is not a trivial exercise, and weights may also need to be changed over time (Mead and Riley 1981). However, in the context of a simulation, the simulated trait also can be considered as the total economic value resulting from a linear selection index. We assume that we would observe similar trends for our simulated breeding programs if we were to include multiple traits in an index.

#### 4.2.4 Assumptions about the absence of genotype-by-genotype interaction between the two component crops

In our simulations, we ignored the possible effects of genotype-by-genotype interactions between component crops. In practical intercrop production, these interactions play an important role in determining their productivity, ecosystem service provision and resilience (Dawson et al. 2019a). Although strategies have been outlined through which genetic variants underlying mutualisms between pairs of plant species in natural ecosystems can be characterised, studies reporting genotype-by-genotype interactions are currently relatively scarce (Subrahmaniam et al. 2018). We expect the effect of genotype-by-genotype interactions to be most significant at the start of breeding activities, when material is unadapted to a particular growing system and when they could potentially result in re-ranking of our breeding programs. We expect that through continuous recurrent selection these interactions may become minimal, as the competition component is minimized through continuously improved coexistence between two component crops (Hill 1996).

### 4.3 Conclusions

Our results show that genomic selection shows great promise in breeding crops for intercrop production. We have demonstrated that genomic selection can significantly increase the rate of genetic gain in intercrop breeding. In particular, the DH-GS breeding strategy provides a simple solution to implement genomic selection on top of an existing phenotypic selection breeding program, without major rearrangements and regardless of the genetic correlation between monocrop yield and intercrop yield. Clearly, the practical challenges of the implementation of genomic selection strategies differ between breeding programs, but we believe that our results will aid breeders in optimizing the implementation process. In our further work we are exploring the utility of different design approaches for crop combinations such as finger millet and groundnut that could be optimised as an important intercrop for reaching multiple human and environmental health benefits in East Africa (Dawson et al. 2019b).

## 5 Author Contributions

JB, JMH and RCG conceived and designed the study. JB developed the plant breeding program simulations. DO and HFO provided information on existing breeding programs. JB, CW and IKD wrote the manuscript, with input from all authors. All authors read and approved the final manuscript.

## 6 Acknowledgements

SRUC authors are grateful for Global Challenge Research Funding on orphan crops (project BB/P022537/1: Formulating Value Chains for Orphan Crops in Africa, 2017-2019, Foundation Award for Global Agriculture and Food Systems).

