## SupplementaryMaterial for "Modelling illustrates that genomic selection provides new opportunities for intercrop breeding"

Abstract

Intercrop breeding programs using genomic selection can produce faster genetic gain than intercrop breeding programs using phenotypic selection. Intercropping is an agricultural practice in which two or more component crops are grown together. It can lead to enhanced soil structure and fertility, improved weed suppression, and better control of pests and diseases. Especially in subsistence agriculture, intercropping has great potential to optimise farming and increase profitability. However, breeding for intercrop varieties is complex as it requires simultaneous improvement of two or more component crops that combine well in the field. We hypothesize that genomic selection can significantly simplify and accelerate the process of breeding crops for intercropping. Therefore, we used stochastic simulation to compare four different intercrop breeding programs implementing genomic selection and an intercrop breeding program entirely based on phenotypic selection. We assumed three different levels of genetic correlation between monocrop grain yield and intercrop grain yield to investigate how the different breeding strategies are impacted by this factor. We found that all four simulated breeding programs using genomic selection produced significantly more intercrop genetic gain than the phenotypic selection program regardless of the genetic correlation with monocrop yield. We suggest a genomic selection strategy which combines monocrop and intercrop trait information to predict general intercropping ability to increase selection accuracy in early stages of a breeding program and to minimize the generation interval.

Supplementary Tab. 1 Summary of *per stage* parameters and annual operational costs for four ‘large’ breeding programs. Cost is set at approximately US $1M. The cost breakdown is shown for the Phenotypic selection breeding program (Pheno) and the three Conventional genomic selection breeding programs.

| **Year** | **Stage** | **Reps** | $\boldsymbol{h}^{\boldsymbol{2}}$ | **Pheno** | **Baseline-GS** | **PYT-GS** | **DH-GS** |
| --- | --- | --- | --- | --- | --- | --- | --- |
| 1 | Cross |  |  | 100 x 100 DH | 100 x 94 DH | 100 x 94 DH | 40 x 160 DH |
| 2 | DH | 1 | 0.10 | 10,000 | 9,400 | 9,400 | 6,400 |
| 3 | PYT | 4 | 0.33 | 1,000 | 1,000 | 1,000 | 1,000 |
| 4 | GIA 1 | 4 | 0.33 | 100¹ | 100¹ | 100¹ | 100¹ |
| 5 | GIA 2 | 24 | 0.50 | 25² | 25² | 25² | 25² |
| 6 | SIA 1 | 16 | 0.67 | 25^*^ | 25^*^ | 25^*^ | 25^*^ |
| 7 | SIA 2 | 32 | 0.80 | 5^*^ | 5^*^ | 5^*^ | 5^*^ |
|  |  |  | Cost (US$) | 988,000 | 986,000 | 986,000 | 992,000 |

Baseline-GS, the Baseline genomic selection breeding program; PYT-GS, the Preliminary yield trial genomic selection breeding program; DH-GS, the Doubled haploid genomic selection breeding program; Reps, the effective number of replications (i.e. locations); h^2^, narrow-sense heritability; DH, the doubled haploid stage; PYT, the preliminary yield trial stage; GIA 1 and 2, the general intercropping ability stages 1 and 2; SIA 1 and 2, the specific intercropping ability stages 1 and 2. ¹ denotes testing with one probe variety; ² denotes testing with three probe varieties; ^*^ number of specific intercrop combinations.

Supplementary Tab. 2 Summary of *per stage* parameters and annual operational costs for the ‘large’ Grid genomic selection breeding program (Grid-GS). Cost is set at approximately US $1M.

| **Year** | **Stage** | **Reps** | $\boldsymbol{h}^{\boldsymbol{2}}$ | **Grid-GS** |
| --- | --- | --- | --- | --- |
| 1 | Cross |  |  | 40 x 160 |
| 2 | DH | 1 | 0.10 | 6,400 |
| 3 | Grid | 1 | 0.10 | 3,600/1,000,000^*^ |
| 4 | SIA1 | 16 | 0.67 | 100^*^ |
| 5 | SIA2 | 32 | 0.80 | 15^*^ |
|  |  |  | Cost (US$) | 988,000 |

Reps, the effective number of replications (i.e. locations); $h^{2},$ narrow-sense heritability; DH, the doubled haploid stage; Grid, the Grid stage; SIA 1 and 2, the specific intercropping ability stages 1 and 2; ^*^ number of specific intercrop combinations.

| **Year** | **Stage** | **Reps** | $\boldsymbol{h}^{\boldsymbol{2}}$ | **Pheno** | **Baseline-GS** | **PYT-GS** | **DH-GS** |
| --- | --- | --- | --- | --- | --- | --- | --- |
| 1 | Cross |  |  | 100 x 25 DH | 100 x 23 DH | 100 x 23 DH | 40 x 40 DH |
| 2 | DH | 1 | 0.10 | 2,500 | 2,300 | 2,300 | 1,600 |
| 3 | PYT | 4 | 0.33 | 250 | 250 | 250 | 250 |
| 4 | GIA 1 | 4 | 0.33 | 25¹ | 25¹ | 25¹ | 25¹ |
| 5 | GIA 2 | 24 | 0.50 | 6² | 6² | 6² | 6² |
| 6 | SIA 1 | 16 | 0.67 | 4^*^ | 4^*^ | 4^*^ | 4^*^ |
| 7 | SIA 2 | 32 | 0.80 | 2^*^ | 2^*^ | 2^*^ | 2^*^ |
|  |  |  | Cost (US$) | 245,800 | 241,800 | 241,800 | 246,800 |

Supplementary Tab. 3 Summary of *per* stage parameters and annual operational costs for four ‘small’ breeding programs. Cost it set at approximately US $250K. The cost breakdown is shown for the Phenotypic selection breeding program (Pheno) and the three Conventional genomic selection breeding programs.

Baseline-GS, the Baseline genomic selection breeding program; PYT-GS, the Preliminary yield trial genomic selection breeding program; DH-GS, the Doubled haploid genomic selection breeding program; Reps, the effective number of replications (i.e. locations); $\boldsymbol{h}^{\boldsymbol{2}}\boldsymbol{,}$ narrow-sense heritability; DH, the doubled haploid stage; PYT, the preliminary yield trial stage; GIA 1 and 2, the general intercropping ability stages 1 and 2; SIA 1 and 2, the specific intercropping ability stages 1 and 2. ¹ denotes testing with one probe variety; ² denotes testing with three probe varieties; ^*^ number of specific intercrop combinations.

Supplementary Tab. 4 Summary of *per* stage parameters and annual operational costs for the ‘small’ Grid genomic selection breeding program (Grid-GS). Cost is set at approximately US $250K.

| **Year** | **Stage** | **Reps** | $\boldsymbol{h}^{\boldsymbol{2}}$ | **Grid-GS** |
| --- | --- | --- | --- | --- |
| 1 | Cross |  |  | 40 x 48 |
| 2 | DH | 1 | 0.10 | 1,920 |
| 3 | Grid | 1 | 0.10 | 225/62,500^*^ |
| 4 | SIA1 | 16 | 0.67 | 25^*^ |
| 5 | SIA2 | 32 | 0.80 | 4^*^ |
|  |  |  | Cost (US$) | 248,850 |

Reps, the effective number of replications (i.e. locations); $h^{2},$ narrow-sense heritability; DH, the doubled haploid stage; Grid, the Grid stage; SIA 1 and 2, the specific intercropping ability stages 1 and 2; ^*^ number of specific intercrop combinations.

|  | **Small budget** | | | | | | | | | | | | | | | | |
| --- | --- | --- | --- | --- | --- | --- | --- | --- | --- | --- | --- | --- | --- | --- | --- | --- | --- |
|  | **corr = 0.4** | | | | |  | **corr = 0.7** | | | | |  | **corr = 0.9** | | | | |
|  | **Pheno** | **Base-GS** | **PYT-GS** | **DH-GS** | **Grid-GS** |  | **Pheno** | **Base-GS** | **PYT-GS** | **DH-GS** | **Grid-GS** |  | **Pheno** | **Base-GS** | **PYT-GS** | **DH-GS** | **Grid-GS** |
| **Pheno** |  | 1.33 | 1.94 | 2.13 | 2.37 |  |  | 1.17 | 1.75 | 2.12 | 1.36 |  |  | 1.19 | 1.75 | 2.23 | 1.1 |
| **Base-GS** | 1.17 |  | 1.46 | 1.6 | 1.78 |  | 1.23 |  | 1.5 | 1.82 | 1.16 |  | 1.27 |  | 1.48 | 1.88 | 0.92 |
| **PYT-GS** | 1.87 | 1.59 |  | 1.1 | 1.22 |  | 2.05 | 1.66 |  | 1.21 | 0.78 |  | 2.01 | 1.59 |  | 1.27 | 0.62 |
| **DH-GS** | 1.74 | 1.49 | 0.93 |  | 1.11 |  | 1.94 | 1.57 | 0.95 |  | 0.64 |  | 1.87 | 1.47 | 0.93 |  | 0.49 |
| **Grid-GS** | 2.21 | 1.89 | 1.19 | 1.27 |  |  | 2.17 | 1.76 | 1.06 | 1.12 |  |  | 1.88 | 1.48 | 0.94 | 1.01 |  |
|  | **Medium budget** | | | | | | | | | | | | | | | | |
|  | **corr = 0.4** | | | | |  | **corr = 0.7** | | | | |  | **corr = 0.9** | | | | |
|  | **Pheno** | **Base-GS** | **PYT-GS** | **DH-GS** | **Grid-GS** |  | **Pheno** | **Base-GS** | **PYT-GS** | **DH-GS** | **Grid-GS** |  | **Pheno** | **Base-GS** | **PYT-GS** | **DH-GS** | **Grid-GS** |
| **Pheno** |  | 1.31 | 1.8 | 2.1 | 2.31 |  |  | 1.26 | 1.86 | 2.46 | 1.58 |  |  | 1.27 | 1.9 | 2.54 | 1.26 |
| **Base-GS** | 1.24 |  | 1.38 | 1.61 | 1.77 |  | 1.3 |  | 1.47 | 1.95 | 1.26 |  | 1.33 |  | 1.5 | 2.01 | 1 |
| **PYT-GS** | 2.06 | 1.66 |  | 1.17 | 1.28 |  | 2.02 | 1.56 |  | 1.32 | 0.85 |  | 2.2 | 1.66 |  | 1.34 | 0.67 |
| **DH-GS** | 1.98 | 1.6 | 0.96 |  | 1.1 |  | 2.21 | 1.71 | 1.1 |  | 0.64 |  | 2.45 | 1.84 | 1.11 |  | 0.5 |
| **Grid-GS** | 2.95 | 2.38 | 1.43 | 1.49 |  |  | 2.78 | 2.15 | 1.38 | 1.26 |  |  | 2.59 | 1.95 | 1.18 | 1.06 |  |
|  | **Big budget** | | | | | | | | | | | | | | | | |
|  | **corr = 0.4** | | | | |  | **corr = 0.7** | | | | |  | **corr = 0.9** | | | | |
|  | **Pheno** | **Base-GS** | **PYT-GS** | **DH-GS** | **Grid-GS** |  | **Pheno** | **Base-GS** | **PYT-GS** | **DH-GS** | **Grid-GS** |  | **Pheno** | **Base-GS** | **PYT-GS** | **DH-GS** | **Grid-GS** |
| **Pheno** |  | 1.44 | 1.99 | 2.3 | 2.53 |  |  | 1.35 | 1.96 | 2.62 | 1.81 |  |  | 1.39 | 2.14 | 2.9 | 1.56 |
| **Base-GS** | 1.28 |  | 1.39 | 1.6 | 1.76 |  | 1.32 |  | 1.46 | 1.94 | 1.34 |  | 1.5 |  | 1.54 | 2.09 | 1.12 |
| **PYT-GS** | 2.39 | 1.87 |  | 1.16 | 1.27 |  | 2.29 | 1.74 |  | 1.33 | 0.92 |  | 2.59 | 1.73 |  | 1.36 | 0.73 |
| **DH-GS** | 2.16 | 1.68 | 0.9 |  | 1.1 |  | 2.54 | 1.93 | 1.11 |  | 0.69 |  | 3.02 | 2.02 | 1.17 |  | 0.54 |
| **Grid-GS** | 4.31 | 3.36 | 1.8 | 2 |  |  | 4.12 | 3.13 | 1.8 | 1.62 |  |  | 3.8 | 2.54 | 1.47 | 1.26 |  |

Supplementary Tab. 5: Summary of ratios for intercrop genetic gain (above diagonal) and intercrop genetic variance (below diagonal). Results are presented for all annual operating budgets and genetic correlations scenarios. Comparisons between breeding programs were done with the values from the final year of the future breeding phase (year 20). Example: Comparison of ratios for Pheno and Grid-GS under medium budget at correlation 0.4. Above diagonal value of 2.31 represents the times genetic gain of Grid-GS was higher compared to Pheno and the below diagonal value of 2.95 represents the times genetic variance of Grid-GS was lower compared to Pheno.

corr, genetic correlation; Pheno, the Phenotypic selection breeding program; Base-GS, the Baseline genomic selection breeding program; PYT-GS, the Preliminary yield trial genomic selection breeding program; DH-GS, the Doubled haploid genomic selection breeding program; Grid-GS, the Grid genomic selection breeding program

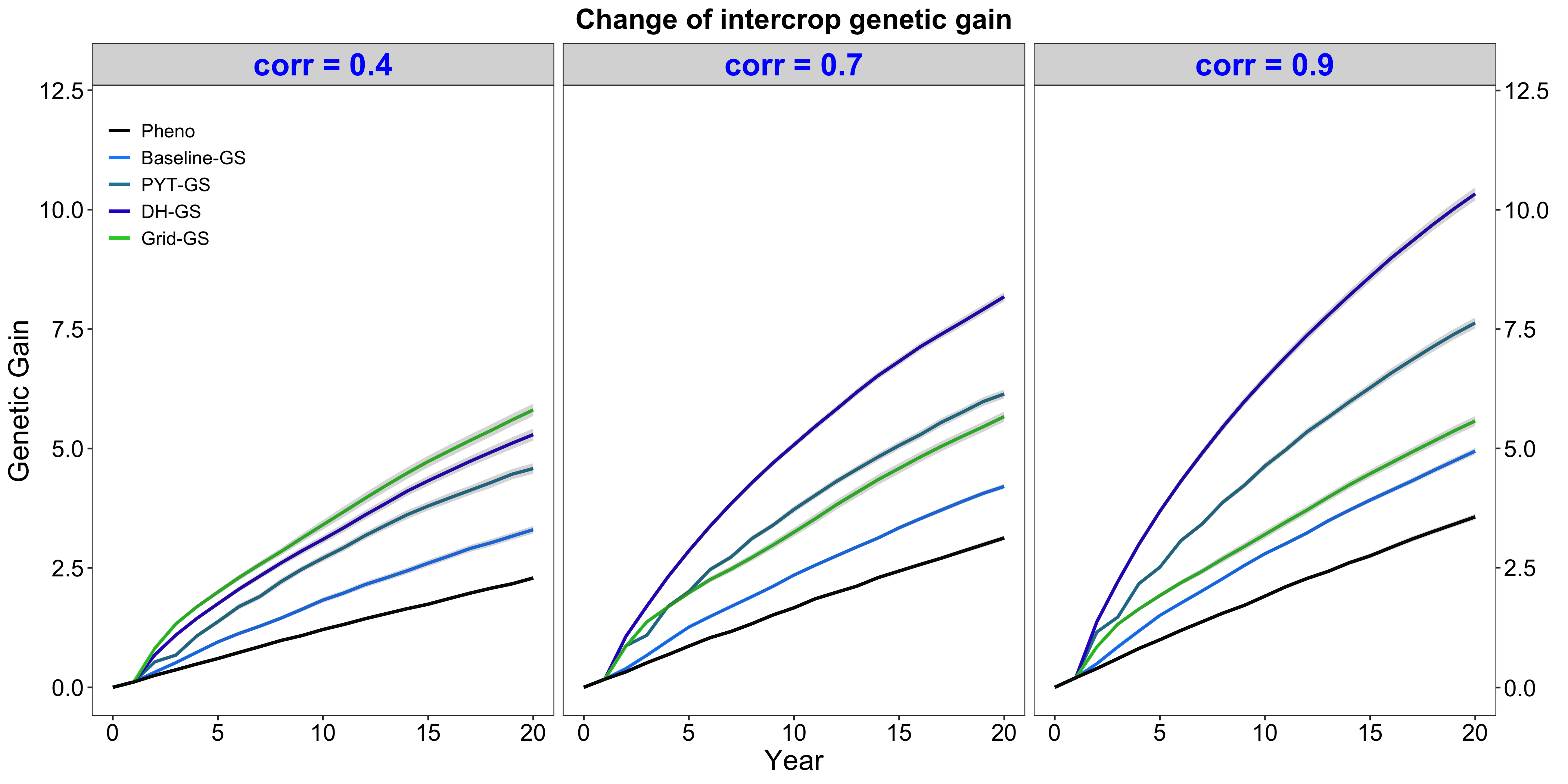

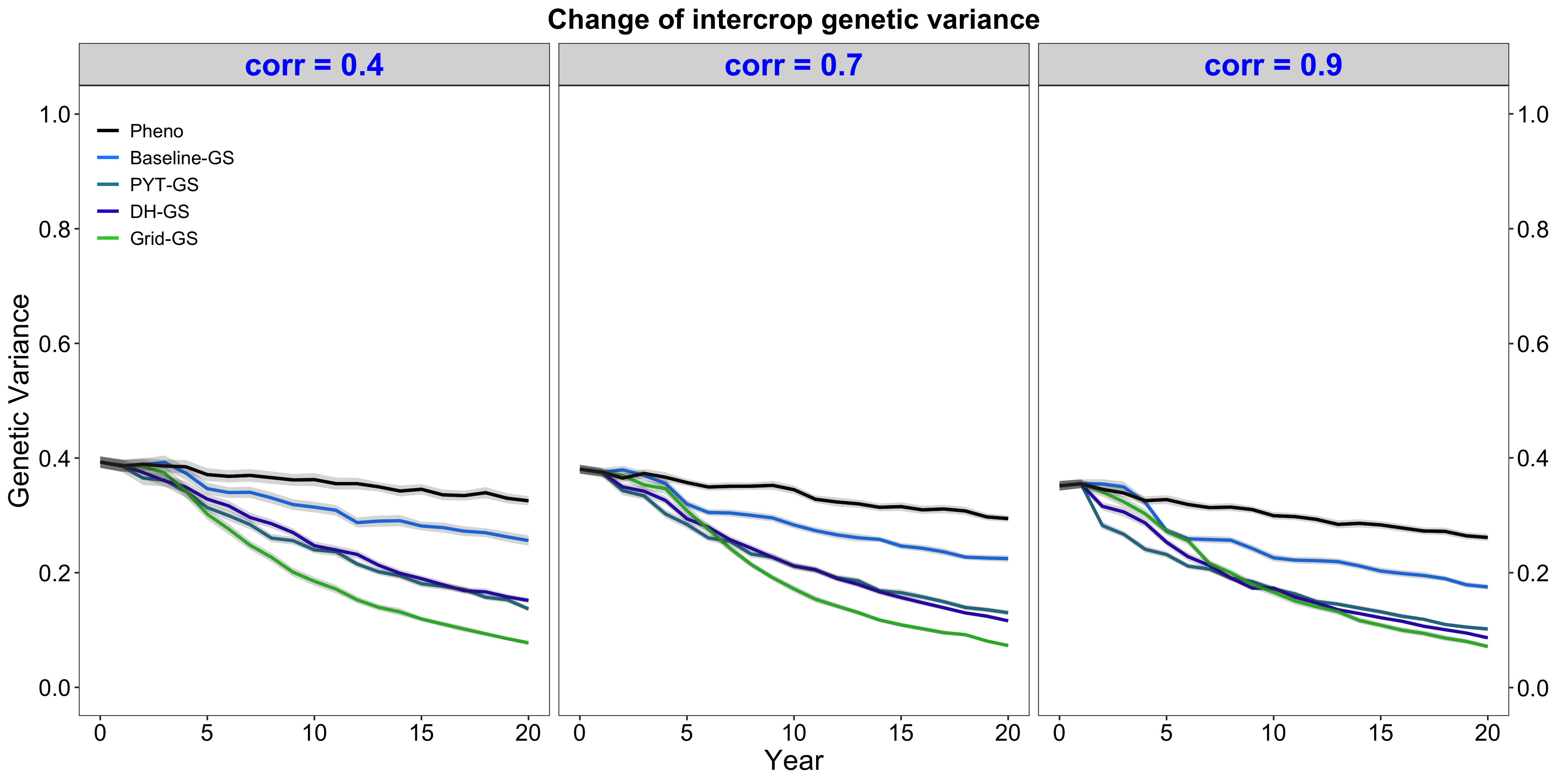

**a)**

**b)**

**c)**

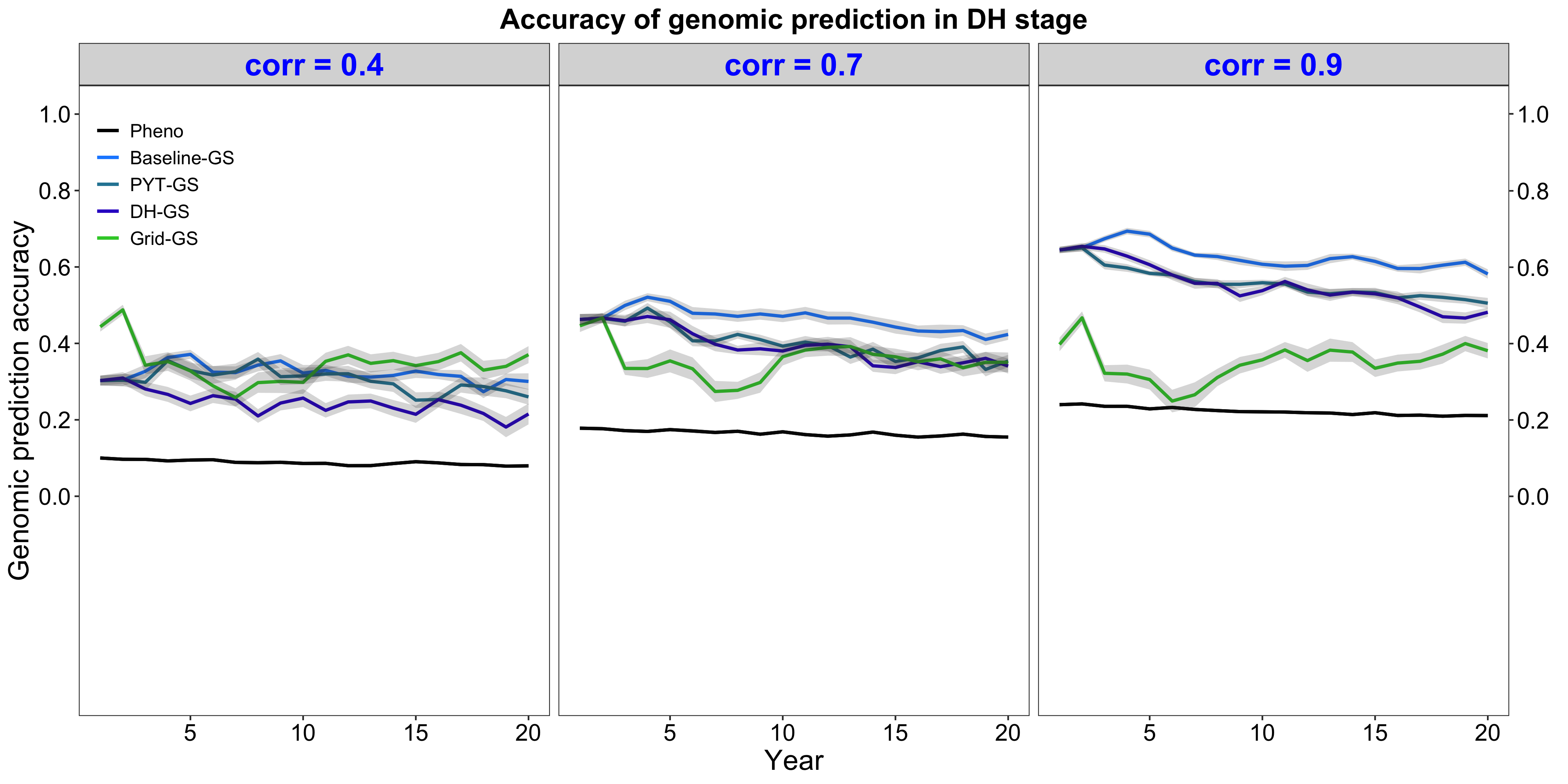

Supplementary Fig. 1. Predicted parameters for ‘large’ breeding programmes. Shown are intercrop genetic gain (a), intercrop genetic variance (b) and genomic prediction accuracy (c) over time for five breeding programs with an annual operating budget of approximately US $1M. Modelling was based on genetic correlations of 0.4, 0.7 and 0.9. Each reported measure is plotted as the mean value for the doubled haploid stage for the entire future breeding phase. The lines within each of the three horizontal panels represent the five breeding programs where each line represents the mean value of the reported measure for the 30 simulated replicates and the shadings visualize conventional error bands. The black line represents the Phenotypic selection breeding program (Pheno), the blue-coloured lines represent the three Conventional genomic selection breeding programs (Baseline-GS, the Baseline genomic selection breeding program; PYT-GS, the Preliminary yield trial genomic selection breeding program; DH-GS, the Doubled haploid genomic selection breeding program) and the green-coloured line represents the Grid genomic selection breeding program (Grid-GS).

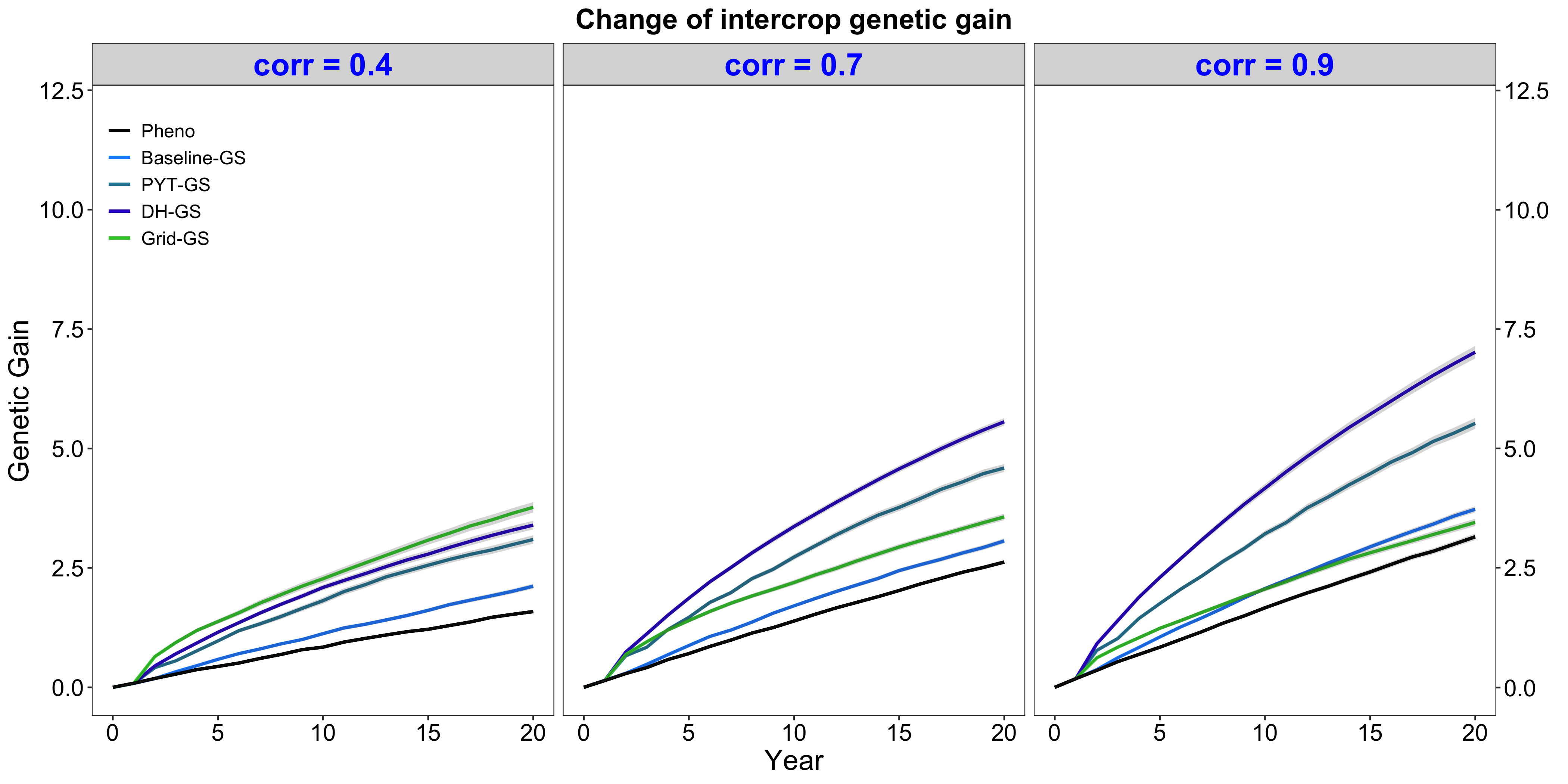

**a)**

**b)**

**c)**

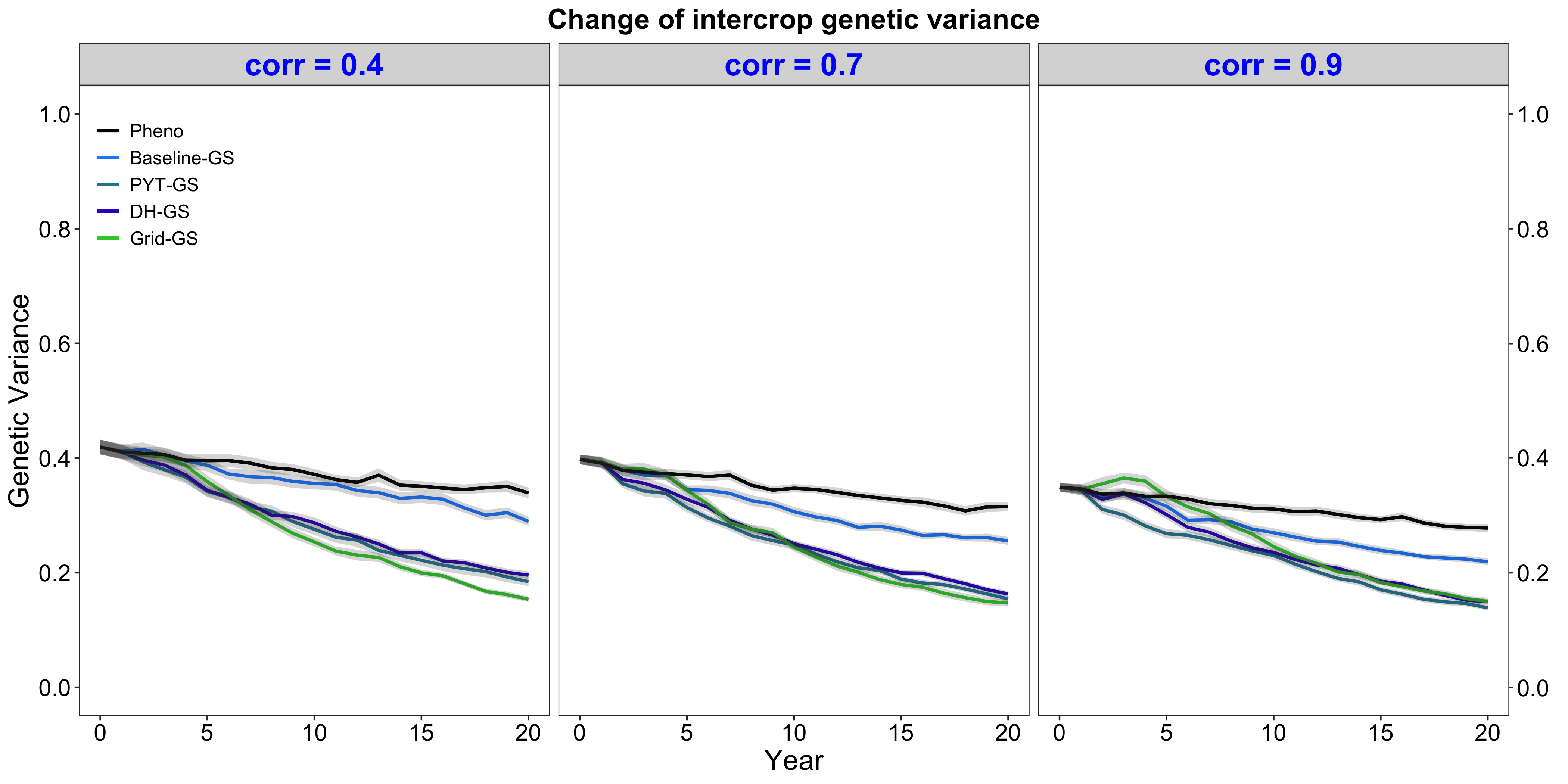

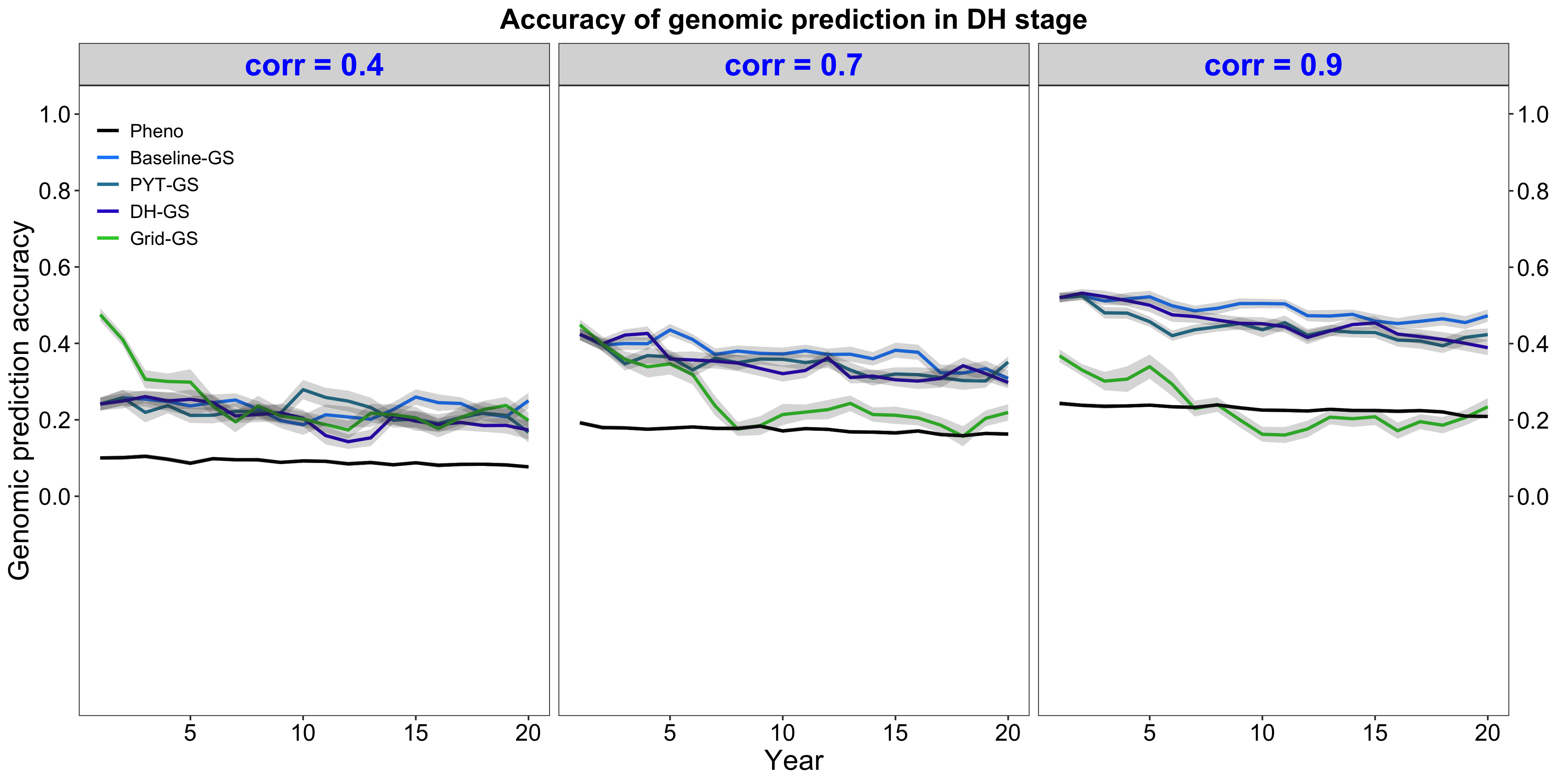

Supplementary Fig. 2. Predicted parameters for ‘small’ breeding programmes. Shown are intercrop genetic gain (a), intercrop genetic variance (b) and genomic prediction accuracy (c) over time for five breeding programs with an annual operating budget of approximately US $250K. Modelling was based on genetic correlations of 0.4, 0.7 and 0.9. Each reported measure is plotted as the mean value for the doubled haploid stage for the entire future breeding phase. The lines within each of the three horizontal panels represent the five breeding programs where each line represents the mean value of the reported measure for the 30 simulated replicates and the shadings visualize conventional error bands. The black line represents the Phenotypic selection breeding program (Pheno), the blue-coloured lines represent the three Conventional genomic selection breeding programs (Baseline-GS, the Baseline genomic selection breeding program; PYT-GS, the Preliminary yield trial genomic selection breeding program; DH-GS, the Doubled haploid genomic selection breeding program) and the green-coloured line represents the Grid genomic selection breeding program (Grid-GS).
